# Dual-feature selectivity enables bidirectional coding in visual cortical neurons

**DOI:** 10.1101/2025.07.16.665209

**Authors:** Nikos Karantzas, Katrin Franke, Konstantin Willeke, Maria Diamantaki, Kandan Ramakrishnan, Hasan Atakan Bedel, Pavithra Elumalai, Kelli Restivo, Paul Fahey, Cate Nealley, Tori Shinn, Gabrielle Garcia, Saumil Patel, Alexander Ecker, Edgar Y. Walker, Emmanouil Froudarakis, Sophia Sanborn, Fabian H. Sinz, Andreas Tolias

## Abstract

Sensory neurons are traditionally viewed as feature detectors that respond with an increase in firing rate to preferred stimuli while remaining unresponsive to others. Here, we identify a dual-feature encoding strategy in macaque visual cortex, wherein many neurons in areas V1 and V4 are selectively tuned to two distinct visual features—one that enhances and one that suppresses activity—around an elevated baseline firing rate. By combining neuronal recordings with functional digital twin models—deep learning-based predictive models of biological neurons—we were able to systematically identify each neuron’s preferred and non-preferred features. These feature pairs served as anchors for a continuous, low-dimensional axis in natural image similarity space, along which neuronal activity varied approximately linearly. Within a single visual area, visual features that strongly or weakly activated individual neurons also had a high probability of modulating the activity of other neurons, suggesting a shared feature selectivity across the population that structures stimulus encoding. We show that this encoding strategy is conserved across species, present in both primary and lateral visual areas of mouse cortex. Dual-feature selectivity is consistent with recent anatomical evidence for feature-specific inhibitory connectivity, complementing the feature-detector principle through circuit mechanisms in which selective excitation and inhibition may together enhance the representational capacity of the neuronal population.

## Introduction

Early sensory physiology established the picture of the single neuron as a feature detector that remains mostly silent until a stimulus in its receptive field resembles a preferred pattern (e.g. Lettvin et al., 1959). This view has intuitive appeal because it suggests that each neuron signals the presence of a specific feature in the sensory world, resulting in a sparse (Barlow, 1961; Field, 1987; Olshausen and Field, 1996) and metabolically efficient code (Levy and Baxter, 1996; Attwell and Laughlin, 2001). For example, in cat and monkey primary visual cortex (V1), a simple cell’s preferred pattern can be described as a specific Gabor function (Hubel and Wiesel, 1962, 1968; Bishop et al., 1973)—a precise combination of orientation, spatial frequency, phase, size, and retinal location. As the similarity of a stimulus to this Gabor increases, the neuron’s firing rate rises, while all dissimilar stimuli evoke little or no activity. Complex cells extend this principle by pooling across spatial phase, yielding phase-invariant orientation tuning (Hubel and Wiesel, 1968; Schiller et al., 1976). This feature detector view has been extrapolated to higher visual areas, where, in the idealized model, a neuron might respond exclusively to a specific face, as in the so-called “Jennifer Aniston neuron” (Quiroga et al., 2005).

Under this framework, lifetime sparsity—the proportion of stimuli that elicit strong firing across a neuron’s entire experience (see also Willmore and Tolhurst, 2001; Willmore et al., 2011)—is determined by the statistics of the preferred feature: neurons tuned to low spatial frequencies tend to fire more frequently because such content is abundant in natural scenes due to their 1*/f* spectral properties (Field, 1987), whereas neurons selective for high frequency edges or specific faces may fire only rarely. While traditionally described in terms of increased firing for preferred features, cortical responses also depend on the suppressive influences that modulate those activations.

Neuronal responses of real cortical neurons, however, reflect a balance of excitatory (i.e. preferred stimulus) and inhibitory inputs (i.e. non-preferred stimulus) within their receptive field. This inhibition is usually considered to be broadly tuned, providing a form of blanket gain control that normalizes neuronal activity across the population (Heeger, 1992; Carandini and Heeger, 2011). A classic example of feature-specific inhibition constitutes cross-orientation inhibition in V1, where responses to a neuron’s preferred orientation are suppressed by the simultaneous presentation of an orthogonal grating (Morrone et al., 1982; Allison et al., 1995; Ferster, 1986). Such examples illustrate that selectivity emerges from the joint action of excitation and inhibition, whose relative balance shapes the final tuning curve. Yet, how this interplay operates in the space of natural images remains poorly understood.

Experimental constraints limit the number of stimuli that can be presented during neuronal recordings, making it challenging to comprehensively map the excitatory and inhibitory influences that determine a neuron’s response across this high-dimensional stimulus space. Resolving this challenge is central to deciphering how neuronal populations transform complex sensory input into meaningful representations.

To overcome this challenge, we leveraged functional digital twin models trained on neuronal recordings from macaque primary (V1) and mid-level (V4) visual cortex, as well as mouse visual cortex. A *digital twin* of a brain area (Walker et al., 2019; Bashivan et al., 2019; Franke et al., 2022; Willeke et al., 2023) can be constructed by training a deep neural network to learn the relationship between stimuli and the resulting neuronal responses. If trained with enough high-quality data, the model can be used to simulate the neuronal response to data never seen by the animal—allowing researchers to scale experiments in silico that would be difficult or impossible in vivo. This approach enabled us to perform a systematic characterization of neuronal selectivity across the full dynamic range of responses to naturalistic images.

We found that many neurons in visual cortex maintain non-zero baseline firing rates, enabling them to exhibit dual-feature selectivity: they respond strongly to preferred features while being systematically suppressed by distinct non-preferred features. We found this bidirectional selectivity not only in primate V1 and V4 but also in mouse V1 as well as lateral visual areas. In addition, we showed that the response function of these neurons reflects a continuum between the preferred and non-preferred features. That is, dual-feature selective neurons exhibit graded responses that vary continuously with perceptual stimulus similarity to their preferred and non-preferred features. By leveraging both excitatory and suppressive selectivity in single cells, this strategy may increase representational capacity by reducing the number of neurons that would be required to encode the equivalent number of features with unidirectional neurons (see section “Balancing sparsity and capacity? A hypothesized role for dual-feature selectivity” in Discussion).

Our results, particularly the observation that non-sparse neurons in visual cortex exhibit feature-selective suppression, align with recent connectomic studies in mice and flies showing specific inhibitory connectivity, where distinct interneuron types target defined excitatory populations (Schneider-Mizell et al., 2025; Matsliah et al., 2024). This contrasts with blanket inhibition, in which inhibitory neurons provide non-selective input that broadly suppresses local excitatory neurons regardless of their feature selectivity (e.g. Heeger, 1992). Such structured inhibitory connectivity suggests that inhibition contributes actively to normalization and selectivity within cortical circuits (see also Sebastian Seung, 2024), thereby complementing the feature-detector principle through mechanisms operating under different circuit constraints to achieve efficient and interpretable neuronal codes.

## Results

### A continuum of response sparseness in macaque V1 and V4

We recorded spiking activity from macaque areas V1 and V4 using linear silicon probes while animals viewed a diverse set of naturalistic images. To align the visual stimulus with the recorded neurons’ receptive fields, we first mapped the population receptive field using sparse noise stimuli while the monkey was fixating on the center of the screen. Probe insertions were targeted orthogonal to the cortical surface, such that neurons sampled along the probe depth share overlapping receptive fields, allowing a single stimulus configuration to adequately drive the entire recorded population. For V1 recordings, the monkey maintained central fixation and we positioned grayscale stimuli (6.7*×*6.7 degrees of visual angle) such that we obtained full receptive field coverage (Fig. 1a left). For V4, we located the fixation dot that centered the population receptive field on the display (Fig. 1a right) and showed full-field RGB images (30*×*16.8 degrees of visual angle). Stimulus sizes were chosen to broadly cover receptive fields across the sampled neural population, including neurons with some spatial scatter around the probe axis (see section “Technical challenges and limitations” in Discussion).

**Fig. 1.**
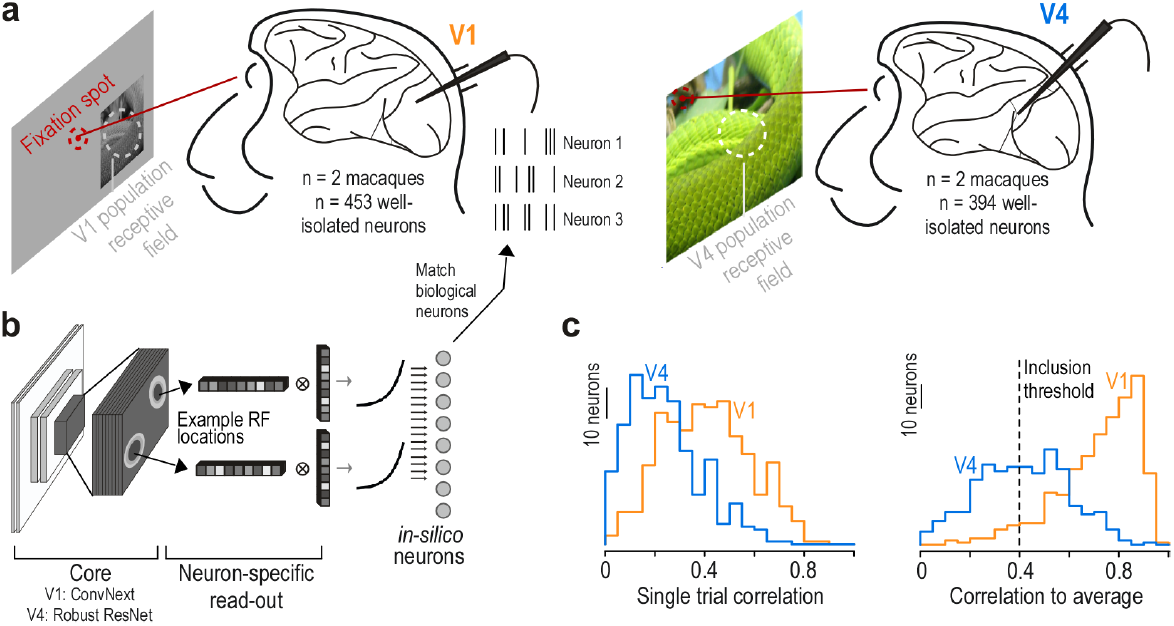
Experimental approach. **a:** Experimental recordings from macaque visual areas V1 (453 neurons from 2 monkeys) and V4 (394 neurons from 2 monkeys) during presentation of natural images. Monkeys fixated on the screen (fixation spot shown in red). The stimulus was centered on the population receptive field of neurons recorded that day, indicated by the white circle. **b:** Architecture of functional “digital twin” models, consisting of a core shared across neurons (V1: first layer of Conv Next; V4: layer 3 of Robust ResNet50) and a neuron-specific readout. This design creates in silico neurons that model the response properties of individual biological neurons, with example receptive field locations of two neurons illustrated. **c:** Prediction accuracy of the model on test images not used during training, for both V1 (orange, *n* = 75 test images) and V4 (blue, *n* = 150 test images). Left panel shows distribution of single trial correlations, and right panel displays correlation to the average response across stimulus repeats. The dotted line indicates an inclusion threshold of 0.4. For further analysis, we only included neurons with a correlation to average above that threshold (*n* = 443 (98% of total) neurons for V1, *n* = 205 (52% of total) neurons for V4).

Monkeys were trained to maintain fixation during 2.1-second trials while we presented 15 images per trial. Each session included between 7, 500 and 15, 000 unique naturalistic images, as well as a smaller set of repeated images to assess response reliability. The V1 dataset was obtained from a previously published study (Cadena et al., 2023), while we collected new data from V4 for this work. After spike sorting, we analyzed data from 453 V1 neurons (*n* = 2 animals) and 394 V4 neurons (*n* = 2 animals).

To characterize each neuron’s stimulus-response function, we trained functional *digital twin* models (Walker et al., 2019; Bashivan et al., 2019; Franke et al., 2022; Willeke et al., 2023). Each model combined a shared convolutional neural network (CNN) core pretrained on image classification (LeCun et al., 2015) with neuron-specific readout layers (Fig. 1b). Our modeling approach leveraged prior findings (e.g. Cadena et al., 2023) that different stages in the visual hierarchy align with different deep neural network layers—early layers with primary visual cortex and deeper layers with intermediate and higher-order areas. Accordingly, we used the first layer of a ConvNeXt architecture for V1 neurons, fine-tuning the convolutional core on the recorded responses and learning neuron-specific readouts (see also Fu et al., 2024). For V4 neurons, we employed a pretrained ResNet50 model as a fixed core and trained a linear readout for each neuron on top of layer 3 of this representation (Willeke et al., 2023).

Model performance was evaluated on test images not used during training (*n* = 75 test images for V1, *n* = 150 test images for V4), using the correlation between predicted and observed responses averaged across stimulus repetitions (Fig. 1c). For further analysis, we included only neurons with correlation-to-average above 0.4, yielding high-confidence *digital twins* for 98% of V1 neurons (*n* = 443) and 52% of V4 neurons (*n* = 205). The lower proportion of high-confidence in-silico neurons in V4 likely reflects the greater complexity of V4 tuning compared to V1, as well as missing contextual information such as image surrounds and sequential image context—factors we discuss in detail below (see section “Technical challenges and limitations” in Discussion). The *digital twins* models enabled us to probe neuronal selectivity across a stimulus space orders of magnitude larger than possible in vivo, where recording time limits the number of stimuli that can be presented per neuron.

To assess response selectivity across natural images, we quantified each neuron’s lifetime sparsity (Willmore and Tolhurst, 2001), which measures how selectively a neuron responds—indicating whether it is activated by many stimuli or only by a few. Using the *digital twin* models, we predicted responses to 1.2 million natural images, generated activation curves by sorting predicted responses in ascending order, and computed their skewness (Fig. 2a,b). Higher skewness reflects one-sided asymmetry in the data and thus indicates sparser response profiles with few stimuli eliciting strong activity. Predicted activity was expressed relative to each neuron’s baseline, defined as the mean response during the 300 ms fixation window immediately preceding stimulus onset, when a uniform gray screen was presented.

**Fig. 2.**
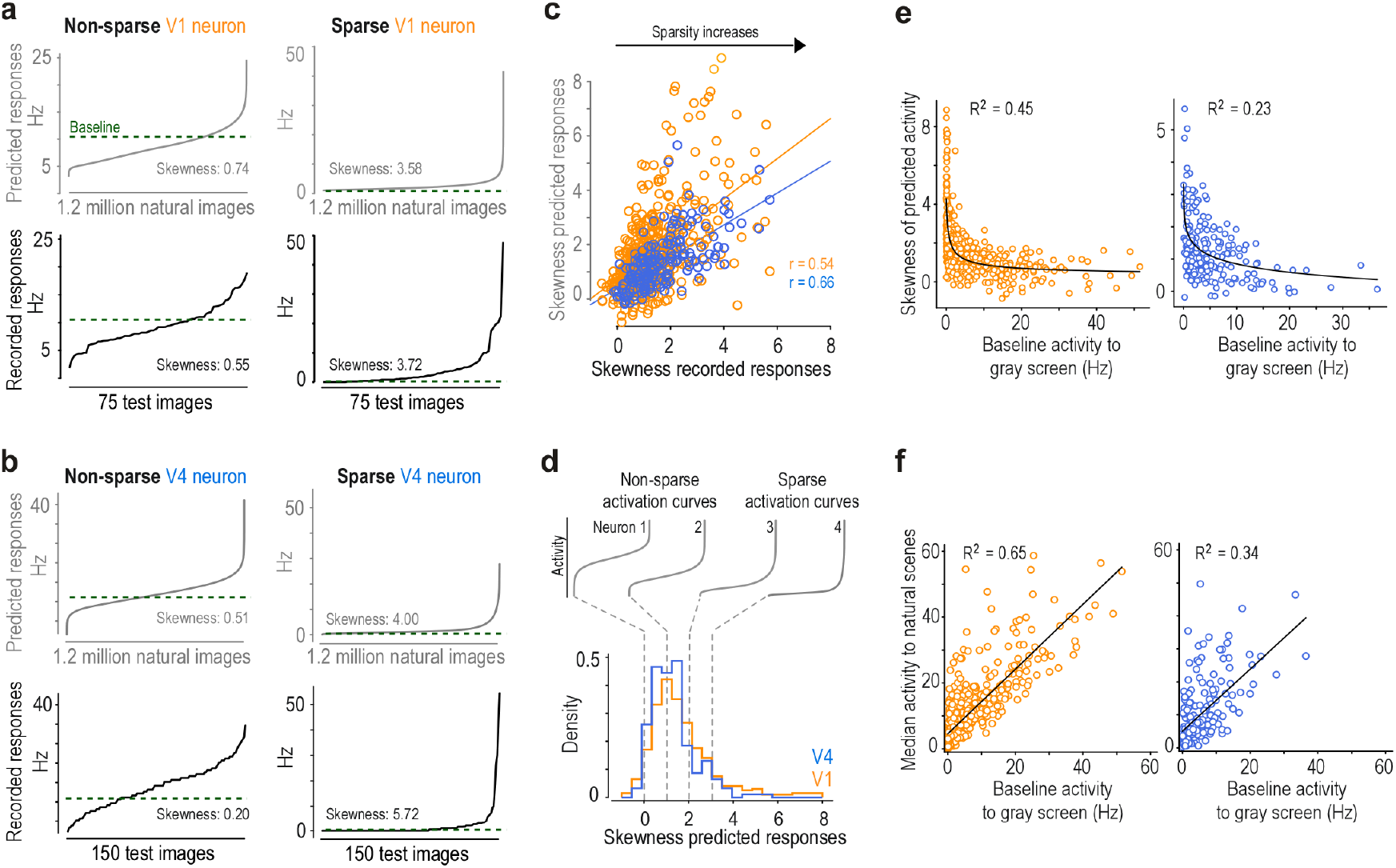
Continuum of single neuron response sparseness in early and mid-level macaque visual cortex. **a**, Response profiles of non-sparse (left) and sparse (right) V1 neurons. Curves display neuronal activity sorted from lowest to highest response, derived from model predictions across 1.2 million ImageNet images (gray, top row) and recorded responses to 75 test images (black, bottom row), averaged over stimulus repeats. We used responses to test images to obtain mean stimulus driven activity and average out stimulus-unrelated signals. Skewness values quantify lifetime sparsity, with higher values indicating neurons that respond selectively to fewer stimuli while remaining silent to most others. Green dotted lines indicate recorded baseline firing rate (Hz) during grey screen presentation prior to stimulus onset. **b**, Comparable response profiles for representative V4 neurons, demonstrating similar variations in lifetime sparsity in this higher-order visual area, with some neurons showing broadly tuned responses and others exhibiting highly selective activation patterns. **c**, Correlation analysis between prediction-based and recording-based skewness values for qualifying V1 (orange, *n* = 443) and V4 (blue, *n* = 205) neurons. The strong correlation (*r* = 0.54 for V1 and *r* = 0.66 for V4, both *p <* 0.001) validates that our model accurately captures intrinsic sparsity characteristics across natural scenes, supporting the use of model-predicted responses for systematic in silico analyses across large image datasets. **d**, Population-level distribution of lifetime sparsity across V1 (orange) and V4 (blue) neuronal populations (V1: *n* = 443 neurons, V4: *n* = 205 neurons), revealing a continuous spectrum rather than discrete categories. Representative activation curves above illustrate how response profiles change along this continuum. Neurons with skewness below 2.0 are defined as non-sparse, though this threshold represents a point along a gradual transition. This distribution highlights the functional diversity within each visual area. **e**, Baseline firing rate extracted from a 300 ms fixation window before stimulus onset plotted versus skewness of predicted responses. V1 (orange): *n* = 443 neurons, V4 (blue): *n* = 205 neurons. *R*^2^ from exponential fit. Neurons with low baseline firing rates exhibit variable skewness that likely reflects the prevalence of their preferred features in natural scenes—rare features produce highly skewed responses while common features yield more symmetric distributions. **f**, Median predicted activity to natural scenes plotted versus baseline firing rate extracted from a 300 ms fixation window before stimulus onset. V1 (orange): *n* = 443 neurons, V4 (blue): *n* = 205 neurons. *R*^2^ from linear regression.

Neurons in both V1 and V4 exhibited activation curves spanning a continuum from highly sparse to non-sparse (Fig. 2d). Skewness values derived from model predictions correlated strongly with those obtained from *in vivo* responses to repeated test images (Fig. 2c), confirming that our *digital twins* accurately capture neuronal stimulus selectivity. Baseline firing rates correlated negatively with response skewness, indicating that neurons with higher baseline activity tended to be less sparse (Fig. 2e). Consistent with this relationship, baseline activity to a gray screen correlated positively with the median response across natural images (Fig. 2f), showing that neurons with elevated baseline firing exhibit response distributions centered near baseline—fluctuating both above and below it. Together, these relationships demonstrate that non-sparse neurons operate in a more continuous, bidirectional modulation regime, while sparse neurons act as highly selective feature detectors. Lifetime sparsity distributions were similar across V1 and V4 (Fig. 2d), with both areas containing substantial proportions of non-sparse neurons (see also Willmore et al., 2011; Rust and DiCarlo, 2012). Model performance correlated only weakly with response skewness, indicating that our inclusion criterion did not systematically exclude sparse neurons from further analysis (Suppl. Fig. 1).

To facilitate subsequent analyses, we used a model-derived skewness of 2.0 to separate neurons with graded, non-sparse responses from those responding strongly to only a small subset of stimuli (Fig. 2d). We note that the underlying distribution of sparsity is continuous, consistent with recent findings (Gondur et al., 2025), and this threshold is adopted purely for analytical convenience to focus subsequent analyses on neurons with sufficiently graded response distributions; the key findings reported below are not dependent on the exact threshold chosen.

### Identification of most and least activating stimuli of non-sparse macaque V1 and V4 neurons

Traditional approaches to visualizing neuronal selectivity have often focused on the most activating stimuli, providing valuable insights into preferred features. Examining weak or suppressive responses, however, can reveal additional aspects of selectivity, particularly in non-sparse neurons that respond in a graded fashion to most stimuli (see section “The role of suppression in visual cortical tuning: relating our findings to existing work” in Discussion). To test whether such weak responses exhibit systematic structure, we characterized both ends of the activation spectrum by identifying stimuli that maximally and minimally activated each neuron using two complementary methods: gradient-based image synthesis and large-scale image screening. Because weak responses are most informative in neurons with graded activity, the following analyses focused specifically on non-sparse neurons, unless noted otherwise.

Based on previous work (Walker et al., 2019; Bashivan et al., 2019; Willeke et al., 2023), the image synthesis approach used gradient ascent in the *digital twin* models to generate images that maximized model-predicted activity. Here, we extended this approach to also minimize neuronal responses. These synthetic stimuli—termed most exciting inputs (MEIs) and least exciting inputs (LEIs)—emerged through iterative modification of noise images to achieve the desired neuronal responses (Fig. 3a). The screening approach computed model activations over more than one million naturalistic images, ranking responses to identify the most activating images (MAIs) and least activating images (LAIs) for each neuron (Fig. 3b). Crucially, all images—both synthesized and screened—were normalized to identical *ℓ*_2_ norms within each neuron’s receptive field, reducing the influence of overall image energy on response differences. We note, however, that L2 normalization controls for root-mean-squared contrast but does not fully equate effective contrast in nonlinear cells, whose responses depend on the spatial structure of the stimulus beyond its total energy. Residual contrast-dependent effects, particularly in the suppressive regime, cannot be entirely excluded.

**Fig. 3.**
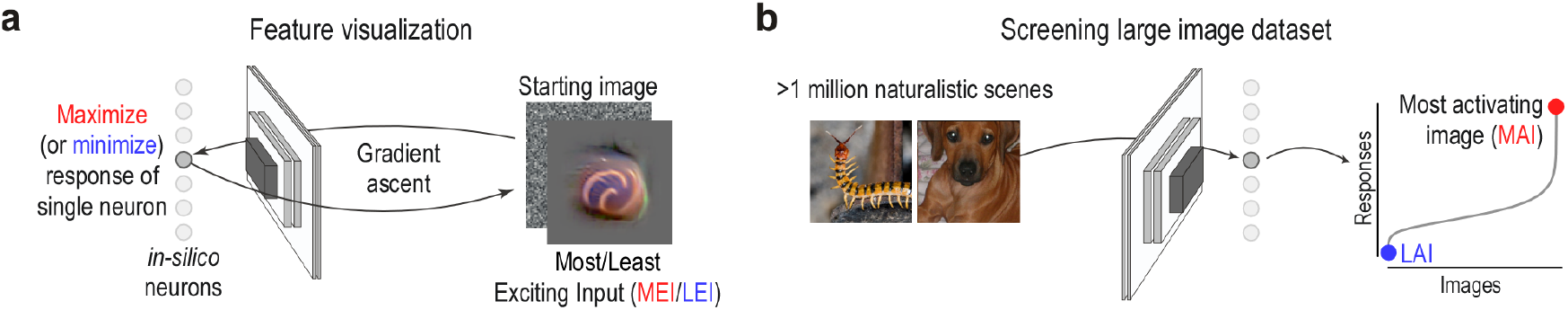
Two complementary methods to study neuronal selectivity: optimization-based feature visualization and large-scale image screening. **a**, Schematic of the feature visualization procedure, in which a starting noise image is iteratively optimized using gradient ascent on a neural predictive model to either maximize or minimize the activity of a single neuron. This yields the most exciting input (MEI) or the least exciting input (LEI), respectively. The process begins with a noise image and updates pixel values to achieve the target activation level. Example shows an LEI for a V4 neuron. **b**, Schematic of the image screening procedure, in which a large dataset of 1.2 million ImageNet images is used to probe neuronal responses. Each image elicits a predicted neuronal response from the model, allowing construction of a response profile across the dataset. Sorting these responses identifies the most activating (MAI) and least activating (LAI) images for each neuron.

In V1, this approach revealed clear structure at both ends of the response spectrum. MEIs and MAIs consistently featured oriented edges and grating-like stimuli spanning various spatial frequencies (Fig. 4, Suppl. Fig. 2), confirming well-established tuning properties of macaque V1 (e.g. Hubel and Wiesel, 1962; Schiller et al., 1976; Fu et al., 2024). Remarkably, LEIs and LAIs exhibited systematic, feature-specific structure: suppression arose not only from orthogonal orientations but also from shifts in orientation, spatial frequency, phase, and texture structure, revealing inhibitory axes that extend beyond classic cross-orientation inhibition (e.g. Morrone et al., 1982, and see section “The role of suppression in visual cortical tuning: relating our findings to existing work” in Discussion). As a control analysis, we simulated idealized simple and complex cells and screened for their least and most activating images (Suppl. Fig. 4). Simple cells showed phase-shifted versions of preferred stimuli as their least-activating images, while complex cells exhibited no coherent patterns in their low-activation regime, with diverse unrelated stimuli eliciting uniformly weak responses—reflecting their pooling mechanisms and nonlinear characteristics. Therefore, the empirical structure we observed—where least activating images differ from the most activating along specific features other than phase—cannot be accounted for by classical simple or complex cell models.

**Fig. 4.**
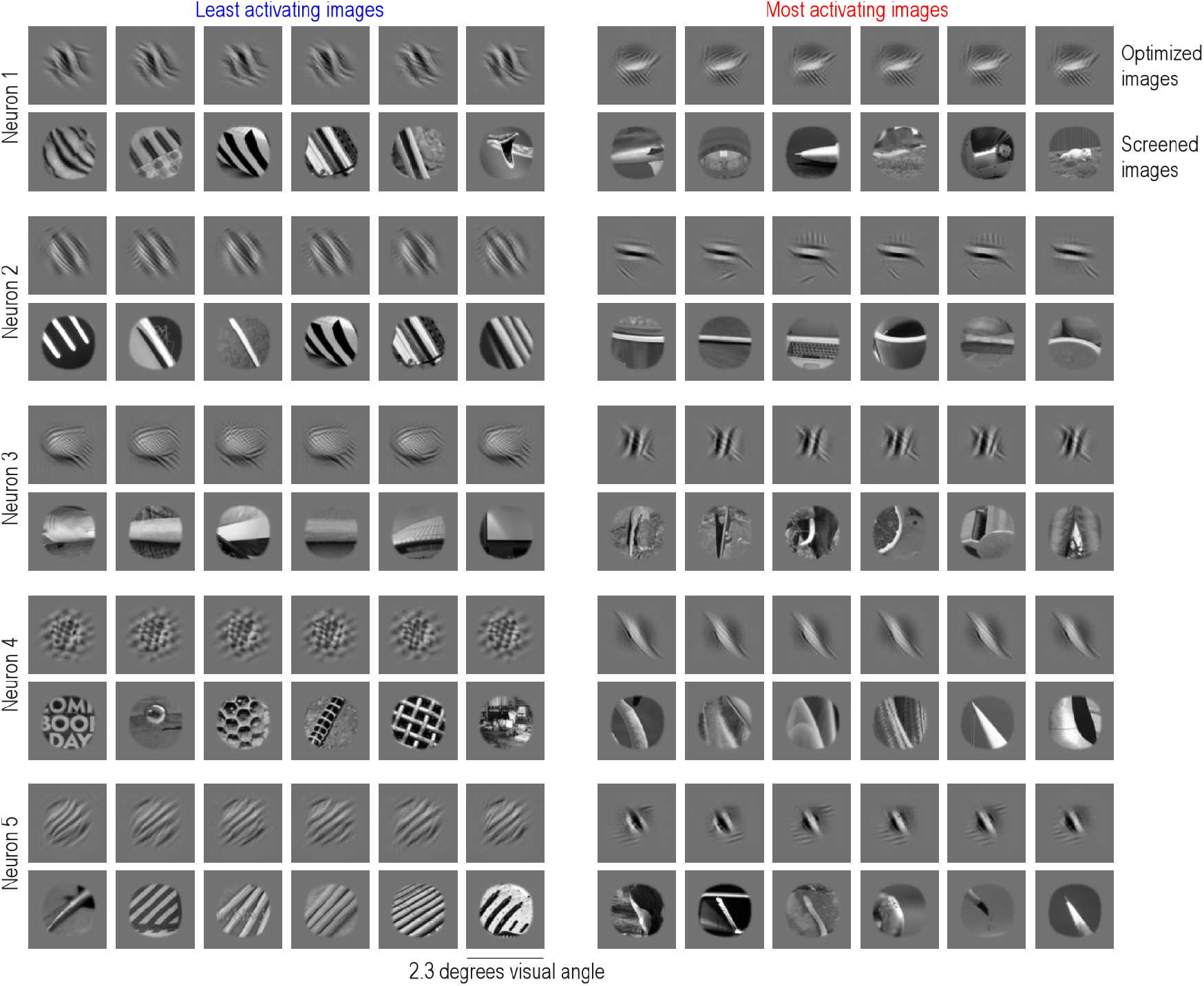
Identification of most and least activating stimuli of macaque V1 neurons. Least (left) and most (right) activating inputs for five example V1 neurons. For each neuron, top row show optimized images starting from different initialization (i.e. noise) seeds (LEIs on the left, MEIs on the right) and bottom row shows the most and least activating images identified through screening 1.2 million ImageNet images (LAIs on the left, MAIs on the right). Images are 2.3 *×* 2.3 degrees visual angle, with each neuron’s receptive field located in the center of the image.

Non-sparse V4 neurons showed similarly structured selectivity at the activation extremes, but for more complex features. Aligning with previous work, their MEIs and MAIs revealed elaborate patterns—including curved contours, textured surfaces, and distinct color combinations (e.g. Willeke et al., 2023; Desimone and Schein, 1987; Pasu-pathy and Connor, 2002; Yamane et al., 2008)—as well as novel motifs such as eye-like configurations and branching structures (Fig. 5, Suppl. Fig. 3). Their corresponding LEIs and LAIs depicted equally coherent feature configurations—alternative contour arrangements, different color combinations, or contrasting texture patterns that consistently suppressed neuronal responses.

**Fig. 5.**
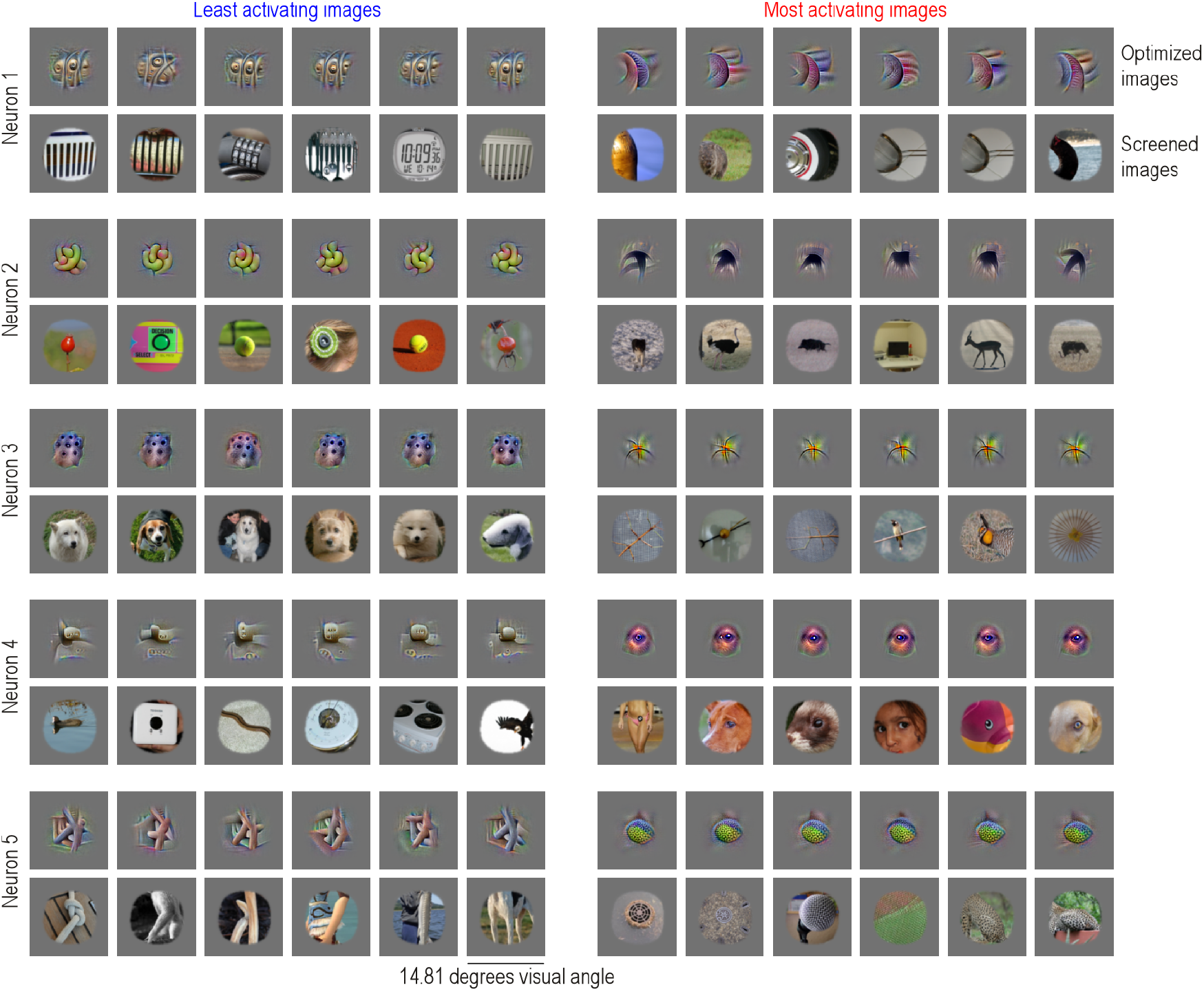
Identification of most and least activating stimuli of macaque V4 neurons. Least (left) and most (right) activating inputs for five example V4 neurons. For each neuron, top row show optimized images starting from different initialization (i.e. noise) seeds (LEIs on the left, MEIs on the right) and bottom row shows the most and least activating images identified through screening 1.2 million ImageNet images (LAIs on the left, MAIs on the right). Images are 14.81 *×* 14.81 degrees visual angle, with each neuron’s receptive field located in the center of the image.

In addition, visual inspection revealed that, in both V1 and V4, the images that most strongly activated a given neuron tended to be perceptually similar to one another, as did the images that elicited the weakest responses. For example, for many V1 neurons, the set of MAIs often shared the same edge orientation but differed in position within the receptive field—consistent with phase invariance in complex cells. In several V4 neurons, the MAIs typically preserved a common global texture or shape, while varying in attributes such as texture phase or color.

To quantify this, we examined the organization of MAIs and LAIs within a perceptual similarity space. We used DreamSim, a model of perceptual similarity fine-tuned to align with human visual judgments (Fu et al., 2023a), and embedded all naturalistic images in this high-dimensional space. For each neuron, we assessed the internal coherence of its preferred (MAIs) and non-preferred (LAIs) image sets by calculating pairwise cosine similarities within the MAIs and separately within the LAIs. We used representations from the penultimate layer of DreamSim, where distances are aligned with human similarity judgments (Fuet al., 2023a). As a baseline, we computed the similarity of the MAIs and the LAIs to randomly selected sets of naturalistic images (Fig. 6a). We found that, for each neuron, the MAIs were more similar to one another than to random images, and the same held for the LAIs (Fig. 6b), resulting in significantly positive d-prime values (Fig. 6c)—a measure of how well each image set could be discriminated from random images. Notably, d-prime values were similar for MAIs and LAIs, demonstrating that low-activating stimuli were just as structured and perceptually coherent as high-activating ones.

**Fig. 6.**
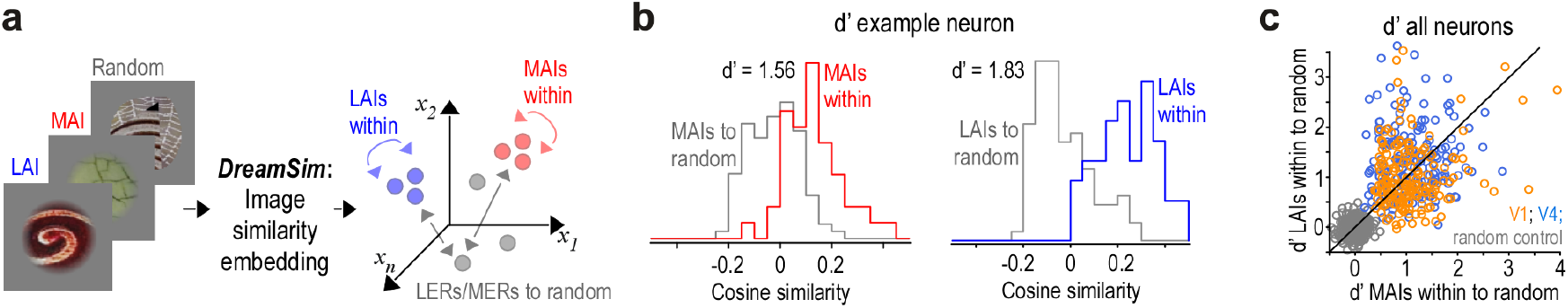
High and low activity reflect structured and perceptually coherent feature combinations. **a**, Schematic illustrating the computation of image similarities. All naturalistic rendered images were embedded into DreamSim, a perceptual similarity space fine-tuned on human judgments. Within this high-dimensional space, we computed cosine similarity among the top 10 most activating images (MAIs) and among the least activating images (LAIs), as well as their similarity to random images. **b**, Distributions of cosine similarity among MAIs (top 10 images, red) and between MAIs and random images (gray). These distributions were used to compute discriminability using the d-prime metric. **c**, d-prime values for MAIs and LAIs across all non-sparse V1 and V4 neurons. Gray bars indicate a control condition comparing similarities between random image sets. Across the population, d-prime values for both MAIs and LAIs were significantly higher in V1 and V4 compared to random image sets (two-sample t-test, *p <* 0.001). After applying false discovery rate correction, all V4 neurons (*n* = 168) retained *p*-values below 0.05, indicating that the likelihood of observing such discriminability by chance was consistently low. In the larger population, *n* = 293 out of *n* = 315 neurons showed significant *p*-values for MAIs, and *n* = 281 neurons showed significant *p*-values for LAIs.

### Verification of most and least activating images of V1 and V4 neurons

Having identified stimuli predicted to generate maximal and minimal neuronal responses, we next validated these model predictions through multiple approaches. Previous studies have confirmed that model-predicted MEIs reliably drive strong responses in both mouse and macaque visual cortex in vivo (Walker et al., 2019; Bashivan et al., 2019; Willeke et al., 2023; Fu et al., 2024). In contrast, whether model-predicted least-activating stimuli suppress neuronal activity remains largely unexplored experimentally (but see Unk, 2025).

For each neuron, we identified the single test-set image predicted by the model to produce the highest response, as well as the one predicted to produce the lowest response, and examined where these images ranked within the neuron’s recorded response distribution over the test set (Fig. 7). For non-sparse V1 and V4 neurons, our models accurately identified both extremes: the image predicted to elicit the strongest response corresponded to a high recorded response, while the predicted least-activating image was associated with responses below baseline, indicating that activity in these neurons was modulated both above and below baseline (Fig. 7a–c, top panels). In contrast, for sparse V1 and V4 neurons, our models reliably predicted the strongest response but performed poorly in identifying weakly activating stimuli (Fig. 7a–c, bottom panels). Because sparse neurons exhibit near-baseline responses for most stimuli, the lower tail of their response distributions has minimal dynamic range. Consequently, the model cannot accurately predict a truly low-activating stimulus, and the recorded responses to predicted least-activating images are broadly distributed across the response range, yielding a nearly uniform distribution.

**Fig. 7.**
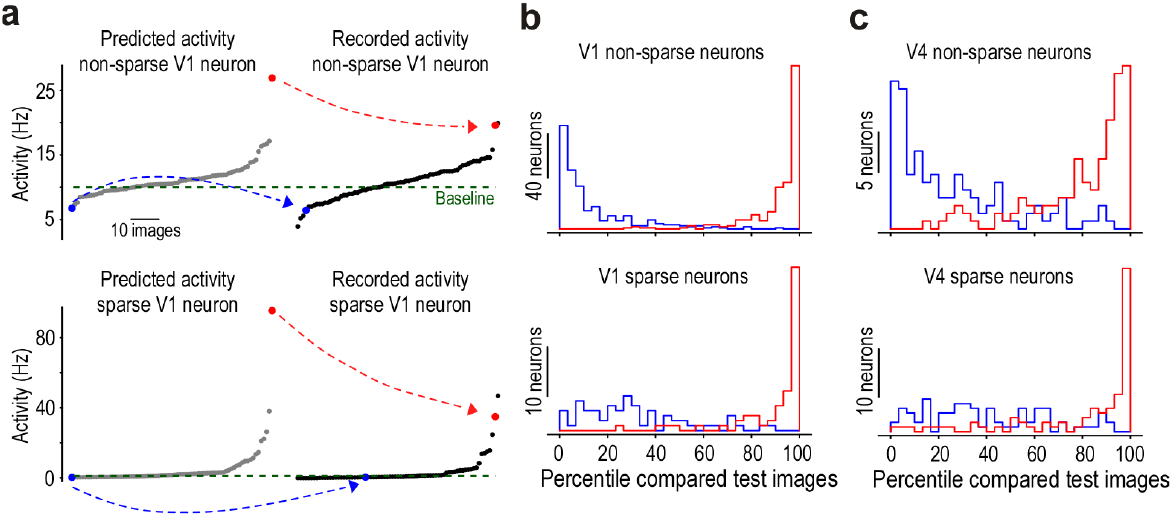
Model predictions accurately identify extreme stimuli in recorded neuronal responses from V1 and V4. **a**, Left: Model-predicted responses to 75 test ImageNet images for an example non-sparse (top) and sparse (bottom) V1 neuron. The predicted least and most activating test images are marked in blue and red, respectively. Right: Actual recorded responses of the same neurons to the same test images, averaged across stimulus repeats and sorted by recorded response magnitude. The blue and red dots indicate the images that the model predicted to elicit the lowest and highest responses. **b**, Distribution of response percentiles in recorded data for the predicted least (blue) and most (red) activating images, shown separately for non-sparse (top) and sparse (bottom) V1 neurons. Note that a non-selective ordering would be expected to yield a uniform distribution, similar to the blue distribution observed here. **c**, Same as (b), but for non-sparse and sparse neurons in area V4.

These results confirm that, for non-sparse neurons, the *digital twin* models capture meaningful stimulus selectivity across both high- and low-activity ranges relative to baseline (Fig. 7a). Although the distribution of response skewness is continuous, a threshold of *skewness* = 2 appears to provide a reasonable separation between neurons with graded, bidirectional modulation around an elevated baseline and those with sparse, predominantly unidirectional responses to a small subset of stimuli (Fig. 2d). For subsequent analyses of least-activating images, we therefore focused on non-sparse neurons with response skewness >2.

Having established that the model predictions align with neuronal responses recorded in vivo, we next asked whether these identified most- and least-activating images reflect genuine neuronal tuning properties rather than artifacts of a particular model implementation. To test this, we evaluated both optimized (i.e. MEIs and LEIs) and screened stimuli (i.e. MAIs and LAIs) using independently trained models with different initializations or architectures. For V1 neurons, we trained an evaluator model with the same ConvNeXt architecture as the original generator model but initialized with independent weights (Fig. 8a). The core was fine-tuned on the same dataset, and neuron-specific readouts were trained from scratch. For V4 neurons, we used a completely different architecture: an attention-based convolutional network trained end-to-end to predict neuronal responses (Pierzchlewicz et al., 2023) (Fig. 8d).

**Fig. 8.**
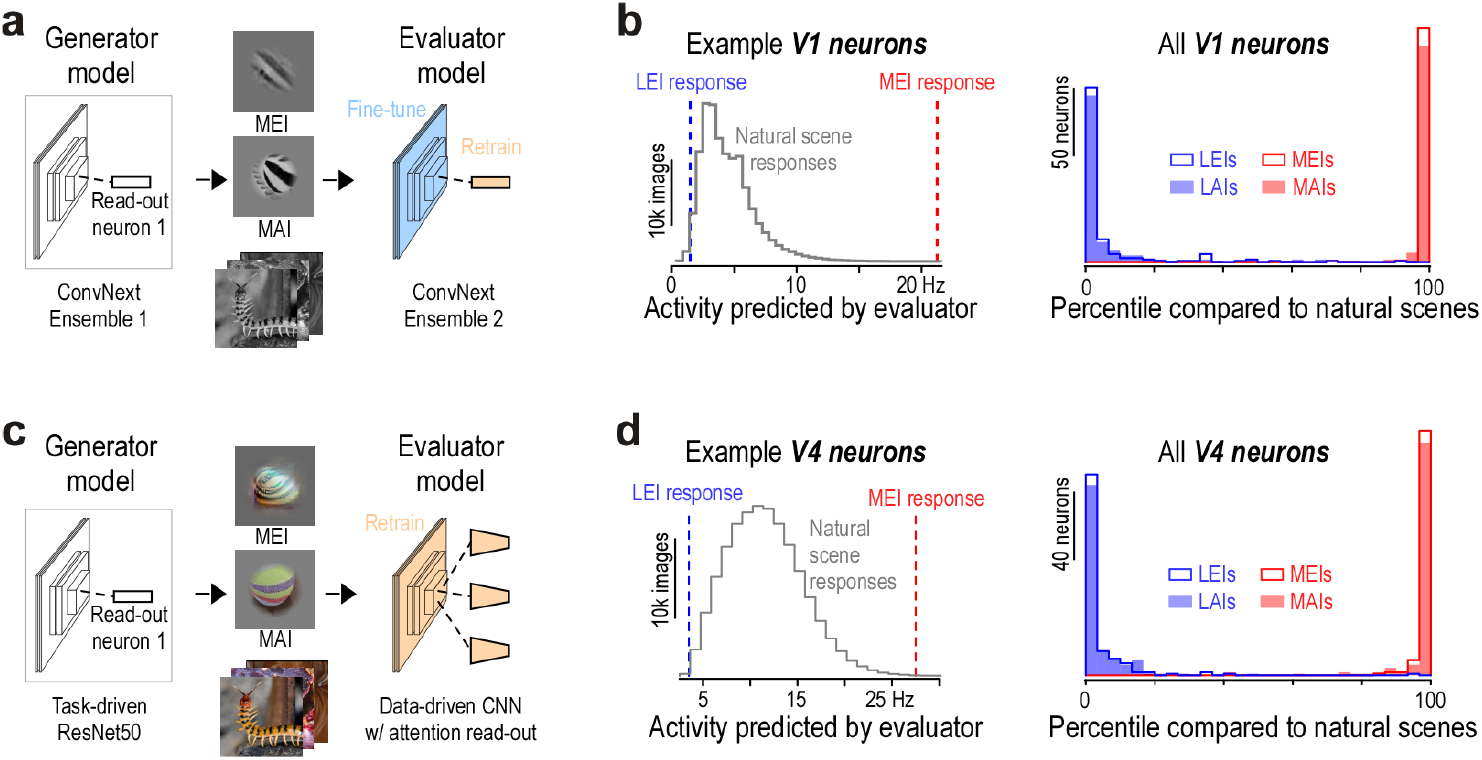
Independent evaluator models confirm the identification of optimal stimuli for V1 and V4 neuronal responses. **a**, Schematic of the verification pipeline. Least and most activating images for V1 neurons were identified using a generator model through both optimization and screening of images. These images were then passed to an independent evaluator model, comprising a separate model ensemble with the same core architecture but a readout trained from scratch. **b**, Example V1 neuron. Left: Distribution of predicted responses to size- and contrast-matched natural images, with the predicted responses to the MEI and LEI (identified by the generator model) highlighted by the evaluator model. Right: Distribution of response percentiles for MEIs, LEIs, MAIs, and LAIs as predicted by the evaluator model, relative to the distribution of natural image responses. MAIs and LAIs were identified based on 200k naturalistic rendered images. **c,d**, As in (a,b), but for V4 neurons. The evaluator model in this case had a distinct architecture and training objective and was trained from scratch on the neural data.

Each evaluator model assessed the MEIs, LEIs, MAIs, and LAIs identified by the generator models. Responses were contextualized against a reference set of 200, 000 naturalistic images that were masked to each neuron’s receptive field and contrast-matched to optimized and screened images. For each neuron, we computed response percentiles—reflecting the proportion of reference images eliciting weaker or stronger model-predicted responses than the identified least- or most-activating images (Fig. 8b,e, left panels). Across both visual areas, MEIs and MAIs consistently ranked in the top percentiles, while LEIs and LAIs fell reliably in the lowest percentiles (Fig. 8b,e, right panels).

These results confirm that for non-sparse neurons, *digital twin* models accurately predict both most- and least-activating images, with stimulus selectivity being robust across independently trained models and architectures, indicating they reflect true neuronal selectivity.

### Feature similarity to most and least activating stimuli jointly shapes non-sparse neuronal responses

Having established that non-sparse neurons are both activated and suppressed around baseline by distinct features, we next asked how these opposing selectivity poles shape their tuning to natural images. Specifically, we sought to understand whether defining both a most- and a least-activating feature imposes structure on how neuronal activity varies across the natural image manifold—that is, whether responses to natural stimuli reflect graded encoding along a continuum between these two poles.

To investigate how neuronal activity varies within the high-dimensional perceptual similarity space, we first constructed a low-dimensional embedding defined by each neuron’s most and least activating features. This allowed us to visualize and quantify how responses scale with perceptual similarity to these features, using a space defined independently of neuronal activity. Specifically, we used DreamSim (Fu et al., 2023a) to embed 200, 000 naturalistic images into a 2-dimensional similarity space for each neuron (Fig. 9a), where each image was assigned coordinates based on its similarity to the MAI (*x*-axis) and LAI (*y*-axis; Fig. 9b). In the following, we focus on V4 neurons because DreamSim captures mid-level visual features like color and texture, which are better represented in V4 than in V1. This alignment is important for relating DreamSim’s similarity space to neuronal activity, as both need to encode similar features. Results for macaque V1 neurons are shown in Suppl. Fig. 5.

**Fig. 9.**
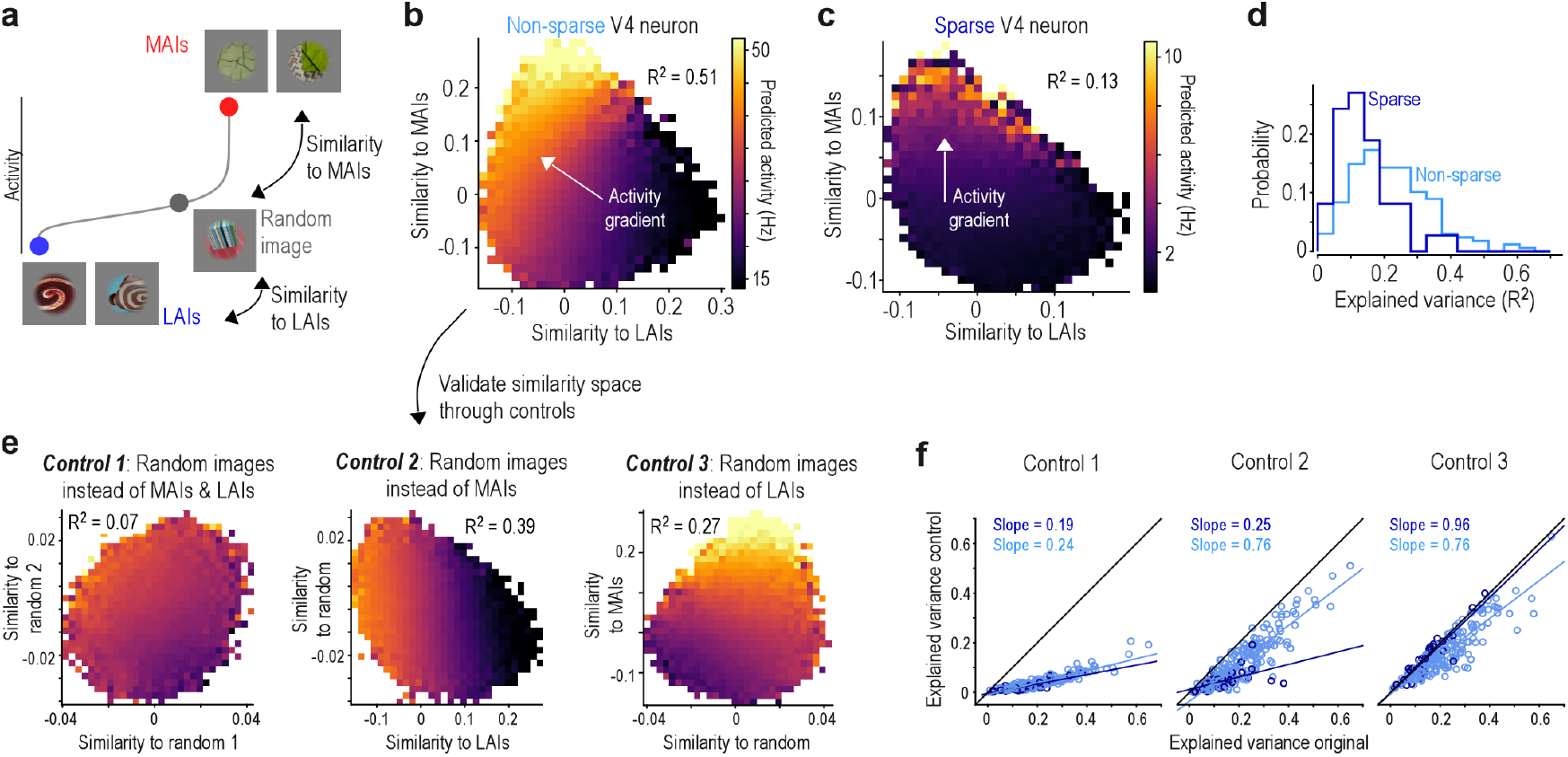
Responses of V4 neurons vary continuously depending on preferred and non-preferred stimuli. **a**, Schematic illustrating construction of the 2D image similarity space for each neuron. Each of *∼*200k naturalistic rendered images was assigned *x*- and *y*-coordinates based on its cosine similarity (computed using DreamSim) to the neuron’s least activating image (LAI, *x*-coordinate) and most activating image (MAI, *y*-coordinate), respectively. MAIs and LAIs were defined from the same image set. **b**, Example 2D similarity space for a non-sparse V4 neuron. Bins are color-coded by the mean predicted neuronal activity of the images within each bin. The arrow denotes the principal activity gradient, and *R*^2^ indicates the variance explained by a linear fit. The color bar spans the 0.1 to 99.9th percentile of neuronal responses across all images. **c**, Same as (b), but for a sparse V4 neuron. Here, the activity gradient aligns primarily along the *y*-axis, indicating stronger modulation by similarity to the MAI than to the LAI. **d**, Variance explained (*R*^2^) by linear regression predicting neuronal activity from the 2D similarity space (as in b,c), shown separately for sparse (dark blue) and non-sparse (light blue) V4 neurons. **e**, Results of three control analyses validating the similarity space for the V4 neuron shown in panel b. Control 1 replaces both MAIs and LAIs with random images. Control 2 replaces MAIs with random images but retains LAIs. Control 3 replaces LAIs with random images but retains MAIs. **f**, Variance explained (*R*^2^) by linear regression for the original analysis (similarity to MAIs and LAIs on *x*- and *y*-axes) and for Control 1 (left), Control 2 (middle), and Control 3 (right). Sparse (dark blue) and non-sparse (light blue) neurons are shown separately. Lines indicate linear regression fits, with slopes reported in each panel.

Visualizing predicted V4 responses across this space revealed that many non-sparse neurons exhibited smooth response gradients along the diagonal, extending from high MAI similarity / low LAI similarity to the opposite corner (Fig. 9b). This gradient indicates that neuronal activity increased with similarity to the MAI and decreased with similarity to the LAI. In contrast, most sparse neurons exhibited less structure, with responses primarily varying with similarity to the MAI (Fig. 9c).

We performed regression to quantify how well the 2-dimensional space explained V4 neuronal activity. The resulting explained variance *R*^2^ was significantly higher for non-sparse neurons (mean=0.23, std=0.12) than for sparse neurons (mean=0.13, std=0.08; Fig. 9d), indicating that the similarity space captures a substantial portion of response variance in the non-sparse population. For V1 neurons, we observed the same pattern of higher explained variance for non-sparse compared to sparse neurons (Suppl. Fig. 5), but overall the explained variance was lower than in V4, despite the *digital twin* model achieving higher prediction accuracy. This is likely because the Dream Sim space is not well aligned with low-level features represented in V1 neurons.

When we repeated the analysis for V4 neurons using randomly selected images instead of MAIs and LAIs (Fig. 9e), the *R*^2^ values were significantly reduced to 0.04 and 0.02 for non-sparse and sparse neurons, respectively (Fig. 9f). This confirms that the observed response gradients depend specifically on the most and least activating images, rather than arising from properties of the similarity space per se. Additionally, for non-sparse neurons, replacing either the MAI or the LAI with random images significantly reduced predictive performance, indicating that both extremes contributed meaningfully to response variation. In contrast, for sparse neurons, only the most activating images carried predictive value; responses remained largely unchanged when the LAIs were replaced by random images.

Together, these findings suggest that the responses of non-sparse neurons are shaped by similarity to both preferred and distinct non-preferred features. Despite the complexity of natural stimuli, V4 responses could be well approximated within a low-dimensional subspace defined by these two feature types, with both contributing meaningfully to response variation.

### A. Shared feature selectivity across the neuronal population

We found that, across neurons within a given visual area, the features that most strongly activate one neuron can resemble those that least activate another (Fig. 10a). Additionally, stimuli that most strongly activate one neuron can also strongly activate others, even when their least-activating images differ (Fig. 10c). These patterns suggest that neurons across the population exhibit shared feature selectivity—that is, they are tuned to a common set of features in perceptual space, which can excite some neurons while suppressing others.

**Fig. 10.**
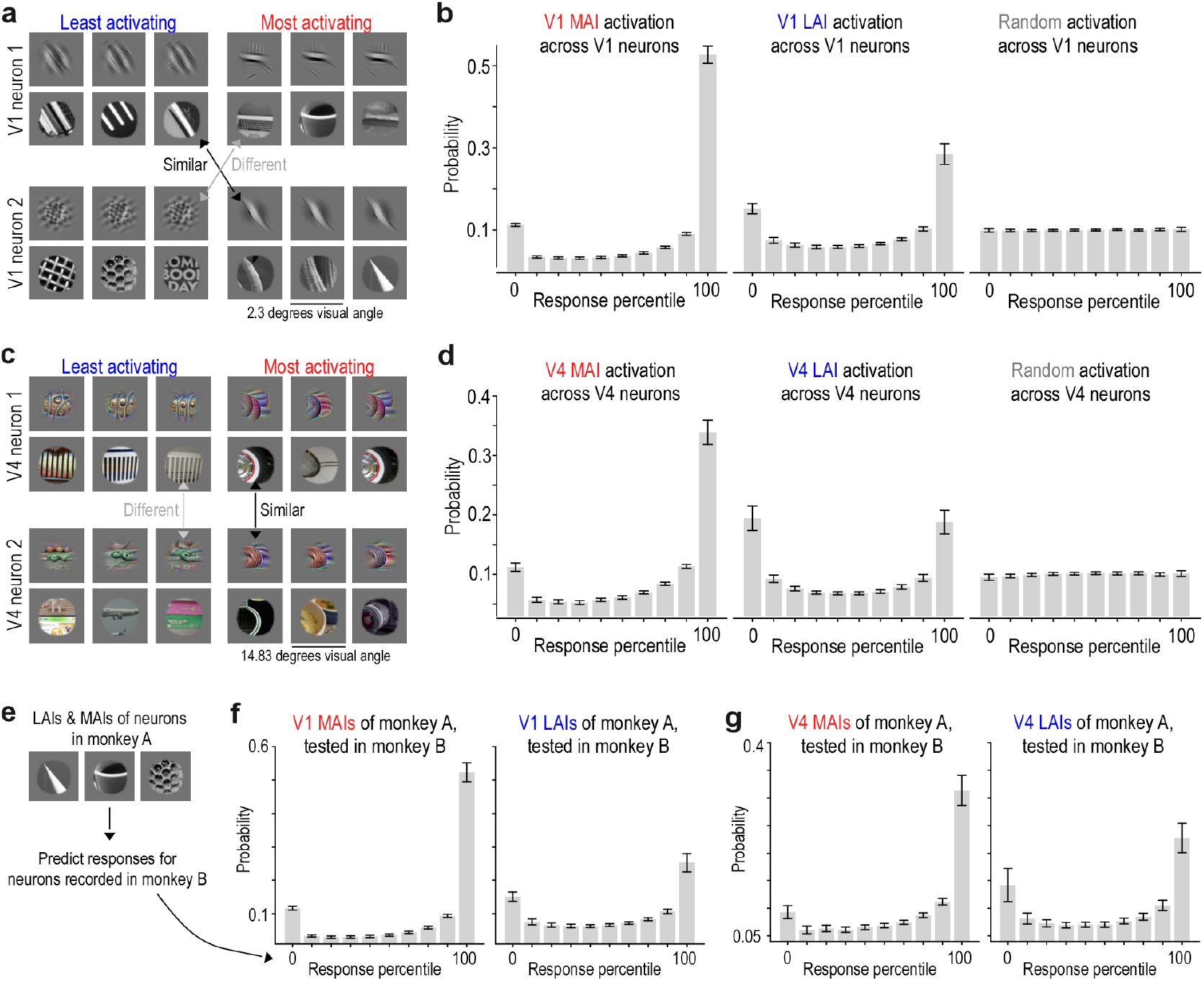
Most and least activating stimuli reveal distributed feature tuning across neuronal populations. **a**, Examples of least activating images (left) and most activating images (right) for two V1 neurons. Arrows indicate the perceptual similarity between MAIs and LAIs shared across both neurons. **b**, Distribution of response percentiles evoked by MAIs, LAIs, and randomly sampled natural images across the V1 neuronal population. The left histogram shows the probability that one neuron’s MAI will elicit a specific response percentile in other neurons. Response percentiles were calculated from each neuron’s responses to 1.2 million ImageNet images. Data represent population means with 99% confidence intervals. **c,d**, Same analyses as in (a,b) applied to neurons recorded in visual area V4. **e**, Schematic illustrating how we used the MAIs and LAIs identified from neurons recorded in one monkey to predict the responses of neurons recorded in another monkey. The resulting response percentiles indicate how these images ranked compared to the predicted responses to 1.2 million ImageNet images, as shown in panels (f,g). **f**, Cross-animal generalization: probability that MAIs (left) and LAIs (right) from V1 neurons in monkey A will evoke specific response percentiles in V1 neurons from monkey B. **g**, Same cross-animal analysis as in (e), applied to V4 neurons.

To test this prediction, we examined how each neuron’s MAIs and LAIs affected the entire population within the same cortical area (Fig. 10b,d). MAIs and LAIs of one neuron showed high probabilities of driving either strong or weak responses in other neurons, while intermediate responses were less common. MAIs elicited right-skewed distributions—with many neurons strongly activated, relatively few weakly activated. LAIs produced bimodal distributions with increased likelihood of both suppression and excitation across the population. This contrasted sharply with randomly selected control images that evoked uniform distributions. This organization extended across individual animals: MAIs and LAIs identified from neurons recorded in one monkey elicited similar activation patterns in neurons recorded from another monkey Fig. 10e-g).

These results support the hypothesis that visual stimuli driving strong or weak responses in individual neurons also modulate responses across the population—exciting some neurons while suppressing others. This pattern reflects a population-level organization of dual-feature selectivity that generalizes across animals.

### DUAL-feature selectivity is present in the mouse visual cortex

To assess whether dual-feature selectivity constitutes a general principle of visual coding across mammals, we extended our analyses to mouse visual cortex. While macaque and mouse visual cortex differ substantially in their functional organization and the complexity of neuronal selectivity (e.g. Fu et al., 2024), we asked whether the broader principle—that non-sparse neurons are jointly defined by distinct excitatory and suppressive feature sets—generalizes across mammalian visual systems. Using Neuropixels probes, we recorded spiking activity from neurons in V1 and two lateral visual areas (lateromedial area (LM) and laterointermediate area (LI)) while head-fixed mice viewed grayscale natural images (Fig.11a). Prior work has shown that LM and LI share functional similarities with the ventral stream in primates (Wang et al., 2012) and are involved in object representation in mice (Froudarakis et al., 2021). Using the recorded data, we trained a *digital twin* model using the Sensorium competition model architecture (Willeke et al., 2022): specifically, a convolutional core shared across neurons combined with neuron-specific readouts, trained end-to-end to predict neuronal responses to natural images (Fig.11b). For further analysis, we only used neurons with a correlation between predicted responses and mean test image responses larger than 0.4. Model performance wascomparable to that obtained in macaque V1, yielding high-confidence digital twins for 92% of neurons across areas, supporting the validity of the subsequent analyses.

**Fig. 11.**
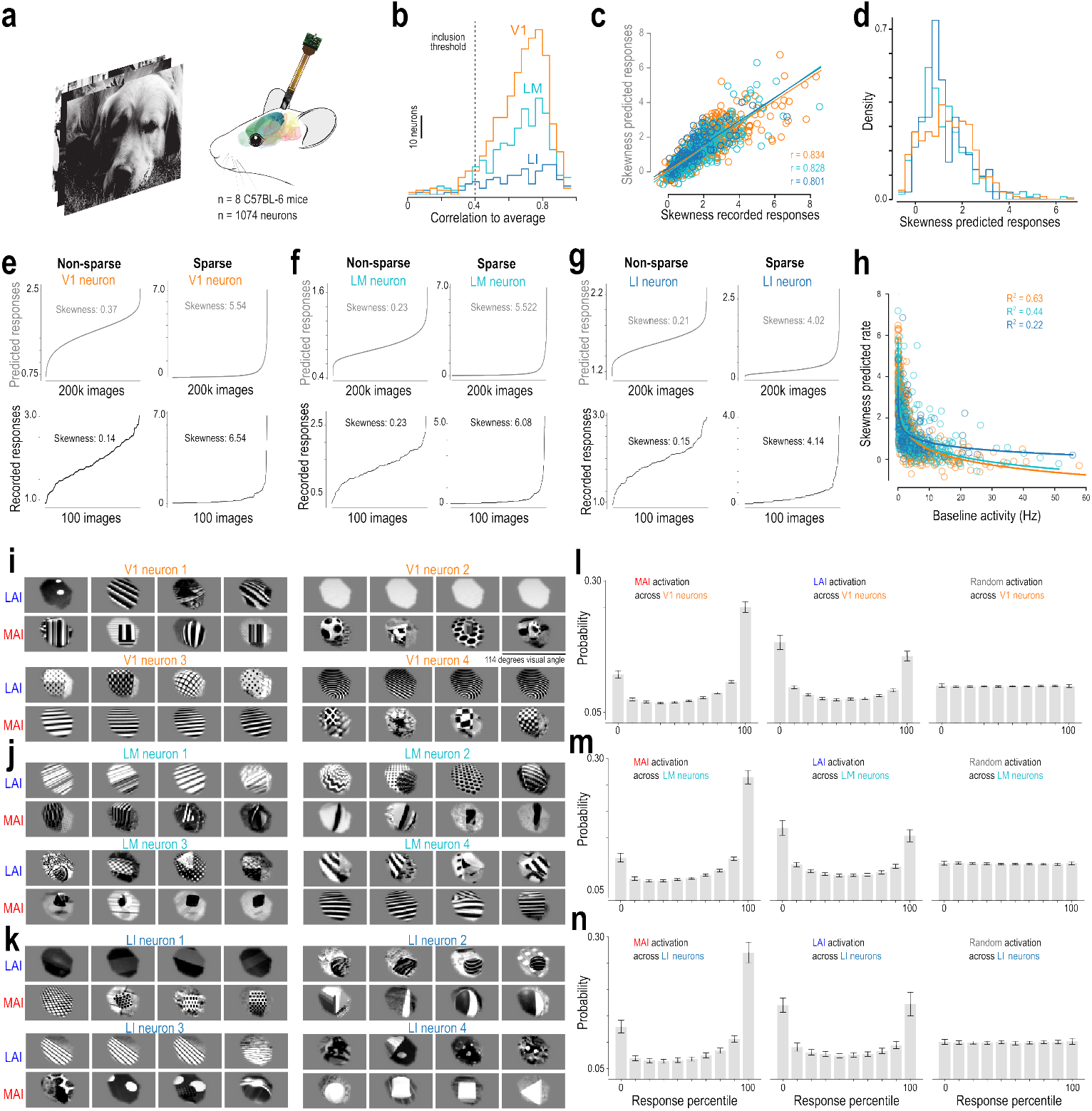
Dual-feature selectivity in the mouse visual cortex. **a**, Experimental recordings from mouse visual areas V1 (598 neurons), LM (350 neurons), and LI (126 neurons) from 8 C57BL-6 mice during presentation of natural images. Mice were head-fixed on a treadmill and passively viewing the stimuli presented on a screen in front of them. **b**, Prediction accuracy of the model on test images not used during training, for V1 (orange), LM (cyan), and LI (blue) as the correlation to the average response across stimulus repeats. The dotted line indicates an inclusion threshold of 0.4. For further analysis, we only included neurons with a correlation to average above that threshold (*n* = 561 neurons for V1, *n* = 325 neurons for LM, *n* = 113 neurons for LI). **c**, Correlation analysis between prediction-based and recording-based skewness values for qualifying V1 (orange, *n* = 561), LM (cyan, *n* = 325), and LI (blue, *n* = 113) neurons. Similar to the primates, there is strong correlation (*r* = 0.834 for V1, *r* = 0.0.828 for LM, and *r* = 0.801 for LI, all *p <* 0.001) indicating that the model accurately captures intrinsic sparsity characteristics across natural scenes. **d**, Population-level distribution of lifetime sparsity across V1 (orange), LM (cyan), and LI (blue) neuronal populations, revealing a continuous spectrum rather than discrete categories. Neurons with skewness below 2.0 are defined as non-sparse. **e**, Response profiles of non-sparse (left) and sparse (right) V1 neurons. Curves display neuronal activity sorted from lowest to highest response, derived from model predictions across 200, 000 ImageNet images (gray, top row) and recorded responses to 100 test images (black, bottom row), averaged over stimulus repeats. Skewness values quantify lifetime sparsity, as in Fig. 2a,b. **f,g**, Same as (e) for LM neurons (f) and LI neurons (g). **h**, Baseline firing rate extracted from a 200 ms window before stimulus onset plotted versus skewness of predicted responses. V1 (orange, *n* = 561), LM (cyan, *n* = 325), LI (blue, *n* = 113) neurons. R^2^ from exponential fit. **i**, Least (blue, LAI) and most (red, MAI) activating images for 4 example non-sparse V1 neurons. LAI and MAI are identified through screening of 200, 000 ImageNet images. Images are 84 x 114 degrees visual angle. **j,k** Same as (i) for LM neurons (j) and LI neurons (k). **l**, Distribution of response percentiles evoked by MAIs, LAIs, and random images across the V1 neuronal population. The left histogram shows the probability that one neuron’s MAI will elicit a specific response percentile in other neurons. Response percentiles were calculated from each neuron’s responses to 200, 000 ImageNet images. Data represent population means with 99% confidence intervals. **m,n** Same analysis as (l) for LM neurons (m) and LI neurons (n).

**Fig. 12.**
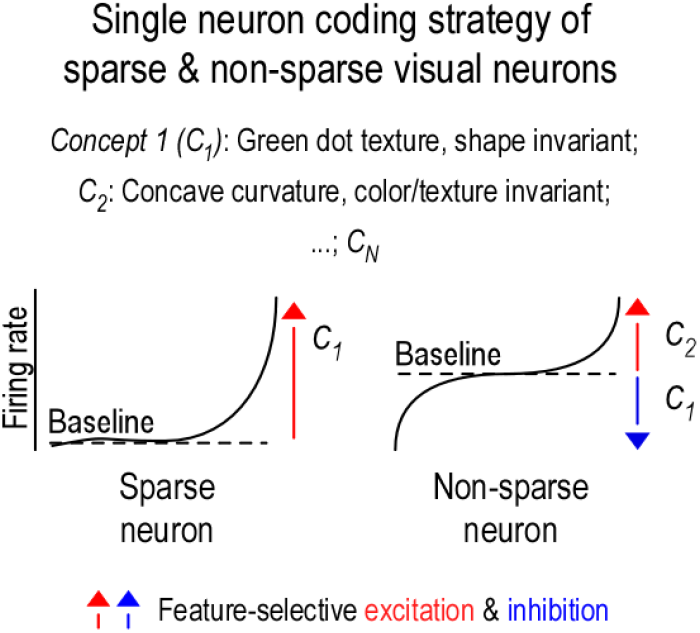
Single neuron coding strategies in sparse and non-sparse visual neurons. This schematic illustrates how sparse and non-sparse neurons exhibit selectivity for distinct *concepts*, where each concept (e.g., *C*_1_, *C*_2_ ) represents a specific combination of latent visual features such as color, shape, and texture—dimensions known to be encoded in area V4. These concepts can be thought of as points in a high-dimensional perceptual space. **Left**: *Sparse neurons* respond selectively and strongly to a single concept (e.g., *C*_1_ : green dot texture) and remain silent for most other stimuli. These neurons exhibit high lifetime sparseness and encode only a narrow portion of stimulus space. **Right**: *Non-sparse neurons* may exhibit *dual-feature selectivity*, characterized by excitation to one concept (e.g., *C*_2_ : concave curvature) and suppression to another (e.g., *C*_1_ ). These neurons tend to maintain non-zero baseline activity and modulate their firing rates bidirectionally, enabling graded responses to a broader range of stimuli. This bidirectional modulation likely reflects feature-selective excitation (red arrows) and inhibition (blue arrows), with firing rates encoding the similarity of each stimulus to both excitatory and suppressive features. Sparse and non-sparse neurons are intermingled within the same cortical area, forming a distributed population code in which different neurons anchor their selectivity to partially overlapping sets of concepts. This organizational motif—shared feature selectivity and bidirectional modulation—appears conserved across visual cortical areas, including earlier regions such as V1, though the underlying concepts differ with the feature space represented at each stage.

Consistent with the macaque data, neurons across all three mouse visual areas exhibited a continuum of life-time sparsity with substantial non-sparse populations, as measured by the skewness of their predicted responses to 200, 000 naturalistic images (Fig.11c,d). The images were masked around the population receptive field and contrast-matched to ensure consistent contrast across images. The skewness of the predicted responses correlated strongly with the skewness of the recorded test responses (Fig.11e-g). Importantly, skewness was negatively correlated with baseline firing rate during gray screen presentation (Fig.11h), suggesting that neurons that were non-sparse during image presentation tended to have higher spontaneous firing rates across areas.

For non-sparse neurons, we next identified LAIs and MAIs by screening a large-scale naturalistic image dataset (Fig.11i-k). As in the monkey, both MAIs and LAIs depicted coherent features. The MAIs of single neurons were perceptually coherent, often featuring specific orientations, textures, or particular visual patterns that drove strong excitation. Similarly, the LAIs showed coherent structure, but these patterns differed systematically from the MAIs along specific dimensions, suggesting that images eliciting minimal responses occupied distinct regions of feature space.

At the population level, the same principle as in the monkey emerged: LAIs and MAIs of a given neuron were more likely to strongly or weakly activate other neurons within the population, compared to randomly selected images which exhibited an approximately uniform likelihood of eliciting responses across the response range (Fig.11l-n).

Together, these findings demonstrate that dual-feature selectivity is present across mouse visual areas, with non-sparse neurons responding to distinct feature sets that drive excitation and suppression respectively. This suggests that the joint definition of neurons by both their preferred and suppressive stimulus features—rather than by excitatory tuning alone—may constitute a general principle of mammalian visual coding. We discuss the potential functional implications of this organization across species below.

## Discussion

Visual cortex has long been understood to represent the visual world through neurons that increase their firing rates in response to specific visual features. Here, we identify an additional organizational principle of single neuron selectivity: many neurons exhibit dual-feature selectivity, responding not only to preferred features but also showing systematic suppression to distinct non-preferred features. This bidirectional selectivity enables individual neurons to continuously encode the contrast between two specific feature combinations in high-dimensional input space. Specifically, the neuron’s firing rate is modulated in a graded fashion, with each rate reflecting the relative distance to the most and least activating stimuli, extending beyond simple feature detection. We find that this principle is present across species (macaque and mouse) and cortical areas (from primary visual cortex to higher-order regions), suggesting it may represent a common computational strategy for single neuron and population coding. These findings indicate that cortical representations integrate both specific excitation and suppression within individual neurons, potentially expanding representational capacity while maintaining interpretable single-neuron responses.

### The role of suppression in visual cortical tuning: relating our findings to existing work

Inhibition plays a fundamental role in sensory processing, with extensive literature documenting its contributions to neuronal encoding (e.g. reviewed in Isaacson and Scanziani, 2011). Here, we focus specifically on visual cortex and suppression originating within the classical receptive field, excluding surround suppression. In primary visual cortex, a classic example is cross-orientation suppression, where a neuron’s response to its preferred orientation is markedly reduced when an orthogonal (non-preferred) grating is presented simultaneously (Bishop et al., 1973; Morrone et al., 1982; Allison et al., 1995; Ferster, 1986; Priebe, 2016; Burr et al., 1981; Hata et al., 1988). Theoretical studies suggest that such inhibitory mechanisms sharpen orientation tuning by suppressing responses to non-optimal stimuli (e.g. Ben-Yishai et al., 1995). Our results reveal that suppression in visual cortex extends beyond well-characterized mechanisms such as cross-orientation inhibition, with single neurons being systematically suppressed by a diverse set of naturalistic features that are distinct from, and seemingly unrelated to (see also Gondur et al., 2025), their excitatory preferences. Using unbiased image synthesis and large-scale screening, we find that macaque V1 neurons’ least activating stimuli are not limited to orthogonal orientations, but comprise specific combinations of orientation, spatial frequency, phase, and size that systematically differ from the features driving maximal activation (Figs. 4, 5). This aligns with previous work demonstrating that suppression extends beyond orthogonal orientations (Ringach et al., 2002; DeAngelis et al., 1992; Burg et al., 2021) and to spatial frequencies outside a neuron’s preferred range (Bauman and Bonds, 1991; De Valois and Tootell, 1983), while going beyond what classical V1 models predict. For simple cells, phase-shifted stimuli are a well-established suppressive axis reflecting linear On-Off subfield structure, and for complex cells, phase pooling yields no coherent suppressive pattern—yet neither model class accounts for the multidimensional suppressive structure we observe (Suppl. Fig. 4). Our unbiased approach reveals that this structure spans simultaneous changes across orientation, spatial frequency, phase, and texture, exceeding what any single known suppressive mechanism predicts.

Importantly, this structured suppression is conserved across species and visual cortical areas. In both mouse primary and lateral visual cortex and macaque V1 and V4, we observe similar feature-specific suppression. Notably, this does not imply that mouse and macaque visual cortex share similar functional organization or equivalent complexity of neuronal selectivity. Rather, within the representational regime of each area—whether mouse V1 or macaque V4 —neurons are organized such that excitatory and suppressive feature sets are jointly structured and distinct, even as the specific features that drive or suppress the neurons differ substantially across species and areas.

Our results support previous evidence for diverse suppressive mechanisms in visual cortex, including cross-orientation inhibition in V2 (Rowekamp and Sharpee, 2017), suppressive subfields in V4 receptive fields (Pollen et al., 2002), suppression by non-optimal stimuli in inferior temporal cortex (Willmore et al., 2011; Rust and DiCarlo, 2012; Miller et al., 1993; Rolls and Tovee, 1995), tuned-suppression to natural images (Tamura et al., 2004), and biased competition among multiple stimuli within receptive fields, modulated by attention (Desimone and Dun-can, 1995; Reynolds et al., 1999). Our work, together with complementary evidence from macaque V4 recordings using multi-unit Utah arrays (Gondur et al., 2025)— which independently demonstrates that V4 neurons exhibit two-tailed response distributions with both preferred and anti-preferred stimuli that are seemingly unrelated in feature space—extends these prior studies by providing a more general framework for how suppression shapes neuronal tuning across the visual hierarchy and species. While prior work has established that inhibition can be structured and feature-selective (see above), our results suggest a broader organizing principle: within each visual area, there exists a set of feature combinations— appropriate to the area’s level of abstraction—rom which individual neurons draw both their excitatory and suppressive preferences. Rather than being idiosyncratic, these preferences are shared across neurons and animals within the same area, such that the features exciting one neuron frequently suppress another. This organization structures the single-neuron code along axes that are meaningful at the population level, giving rise to graded, bidirectional modulation around baseline that reflects a stimulus’s position along feature dimensions collectively relevant to the area’s computational role.

A directly analogous organizational principle has been described in face-selective cortex. Chang & Tsao demonstrated that face identity is encoded through a set of approximately 50 shared axes spanning a low-dimensional face space, with individual neurons responding as linear projections onto these axes (Chang and Tsao, 2017). Rather than representing idiosyncratic faces, face-selective neurons collectively tile a shared manifold in which each axis captures a specific facial dimension. Our results suggest that a related principle may extend beyond the specialized domain of face perception to general visual cortex: each non-sparse neuron anchors a selectivity axis in natural image space—defined by its most and least activating stimuli—and responds in a graded, approximately linear fashion as images vary along this axis (Fig. 9). Where Chang & Tsao characterized axes within the constrained geometry of face images, a central question for future work is whether the selectivity axes of V4 neurons—spanning arbitrary combinations of color, texture, curvature, and object structure—likewise decompose into a compact set of shared primitives reused across the neuronal population. A related geometric organization has been observed in artificial vision systems, where self-supervised networks such as DINO exhibit population-level “concept axes” with antipodal poles encoding opposing semantics (Fel et al., 2025). Our results reveal a distinct principle operating at the single-neuron level: biological axes link structured, non-opponent features within a continuous representational manifold.

### Balancing sparsity and capacity? A hypothesized role for dual-feature selectivity

Dual-feature selectivity arises when neurons exhibit bidirectional modulation to two distinct stimulus features, enabling a coding regime that may balance interpretability with representational capacity. Unlike classical feature detectors, which respond sparsely to a narrow set of stimuli (Barlow, 1961; Field, 1987; Olshausen and Field, 1996), dual-feature selective neurons respond to both excitatory and suppressive inputs, potentially spanning a broader dynamic range and likely supporting higher information-theoretic capacity.

Similar geometric organization has been observed in artificial vision systems, where self-supervised networks such as DINO exhibit population-level “concept axes” with antipodal poles encoding opposing semantics (Fel et al., 2025). Our results reveal a related but distinct principle operating at the single-neuron level: biological axes link structured, non-opponent features within a continuous representational manifold, extending the notion of bidirectional coding to the geometry of individual neurons.

By producing distinct firing rates for excitatory, neutral, and suppressive stimuli, such neurons increase the diversity of their response distributions. Mutual information between stimulus and response increases when the response distribution is both diverse—i.e., has high entropy—and reliable—i.e., exhibits low variability given the same stimulus. Formally, mutual information is defined as *I*(*x*; *r*) = *H*(*r*) − *H*(*r* | *x*), where *H*(*r*) is the entropy of the response distribution and *H*(*r* | *x*) is the conditional entropy reflecting noise or ambiguity in the response. Neurons with bidirectional selectivity can increase *H*(*r*) by utilizing a broader dynamic range—responding with different firing rates to excitatory, neutral, and suppressive stimuli—resulting in a more uniform and distributed response profile. If these response patterns are reliable across repeated presentations of the same stimulus (i.e., *H*(*r* | *x*) remains low), then mutual information is increased. This strategy parallels mixed selectivity in higher cognitive areas, where neurons encode nonlinear combinations of features to support high-dimensional and flexible representations (Rigotti et al., 2013; Fusi et al., 2016).

The benefit of dual-feature coding, however, depends on the alignment between a neuron’s selectivity and the statistics of the input space. If excitatory and suppressive features occur frequently and independently, the neuron can fully exploit its dynamic range, distributing responses more uniformly and maximizing entropy. If the features are strongly correlated or rarely occur in natural scenes, responses become concentrated in a narrow range, limiting entropy and diminishing the coding benefit. Thus, the information-theoretic capacity of a dual-feature neuron is tightly constrained by how well its selectivity structure aligns with the distribution of stimuli it encounters.

Neuronal codes with high entropy—where individual neurons respond to many stimuli with variable firing rates—can enhance representational capacity but come with the trade-off of increased metabolic cost. This is because generating action potentials and driving postsynaptic glutamatergic currents are among the most energy-demanding processes in the brain (Levy and Baxter, 1996; Attwell and Laughlin, 2001). Yet, structured population responses may help offset this cost. We find that the same stimulus can increase firing in some neurons while suppressing others below baseline, which may reduce net activity and maintain population sparseness. The resulting response variance across the population accounts for a previous finding, where stimuli optimized to elicit high activity in one area simultaneously suppress certain neurons, thereby expanding the dynamic range (Tong et al., 2023).

Importantly, single neuron lifetime sparseness and population sparseness, while often correlated (e.g., Froudarakis et al., 2014), capture distinct coding properties. As argued by Willmore and Tolhurst (2001); Willmore et al. (2011), population sparseness can remain high even when individual neurons are broadly tuned—so long as different neurons are active for different stimuli. Therefore, dual-feature selective neurons with high firing rates could be organized as a population to maintain balanced population sparseness, ensuring a broad dynamic range while minimizing metabolic costs.

### Technical challenges and limitations

This study has several important limitations that merit careful consideration. First and foremost, our conclusions rely partially on in silico analyses employing *digital twin* models rather than direct experimental validation. Although digital twin models confer substantial advantages that direct analysis of experimentally recorded responses cannot match— enabling screening of more than one million images per neuron in silico, gradient-based synthesis of stimuli precisely optimized to drive or suppress individual neurons, and cross-model verification of identified selectivity patterns (Fig. 8)—the reliance on model-predicted rather than experimentally measured responses requires careful justification. We maintain confidence in our findings for several reasons. Prior research has consistently demonstrated that digital twin approaches can successfully predict neuronal responses to novel stimuli and identify maximally exciting inputs subsequently verified in vivo (Walker et al., 2019; Bashivan et al., 2019; Franke et al., 2022; Willeke et al., 2023; Fu et al., 2024; Tong et al., 2023). We restricted analyses to neurons exhibiting high predictive accuracy on held-out test data, ensuring models faithfully captured neuronal response functions. We further confirmed that model accuracy remained consistent across both high- and low-activity regimes relative to baseline— though exclusively for non-sparse neurons (cf. Fig. 7). Critically, cross-model verification confirms that identified stimuli reflect genuine neuronal tuning rather than model artifacts—a test that has no analog when working with fixed experimental image sets (cf. Fig. 8).

A related consideration concerns receptive field coverage and its contribution to performance differences between V1 and V4. Although we targeted orthogonal probe insertions and centered stimuli on the population receptive field prior to recording, partial receptive field drive for individual neurons cannot be excluded. Given the larger and more variable receptive fields in V4, this may have contributed to lower model performance there. However, other factors are likely more consequential: we cropped images for computational tractability, potentially losing contextual information that may modulate V4 responses; our models did not account for sequential image context during training; and more recent backbone architectures such as DI-NOv2 (Oquab et al., 2023) would likely improve predictivity. Future work moving towards continuous dynamic models and state-of-the-art architectures (e.g. Willeke et al., 2025) will likely close this performance gap, particularly for higher visual areas where temporal context and largescale image statistics play a more prominent role. Crucially, the goal here was not to maximize predictive performance per se, but to identify response patterns—dual-feature selectivity—that are robust across neurons, areas, and species, and our restriction to high-confidence neurons provides resilience against these limitations.

A further consideration concerns generalizability across individual animals. Both V1 and V4 data were collected from 2 macaques. For V4, digital twin models were fit independently per neuron without sharing information across animals. For V1, while models shared a common backbone fine-tuned on recorded data, readout layers remained neuron-specific, ensuring individual response functions are learned from each neuron’s own data. Critically, extreme image sets identified by the model elicited correspondingly extreme responses in neurons from another animal, confirming that identified selectivity patterns are not idiosyncratic to individual subjects. Equivalent results in mouse visual cortex. Together, these results suggest that dual-feature selectivity is a robust and general property of mammalian visual cortex, rather than an artifact of individual animals, recording sessions, or modeling choices.

Another important consideration concerns the scope of our characterization. While we demonstrate dual-feature selectivity in spiking responses, our analyses do not determine the underlying circuit mechanisms, for example whether the observed suppression arises from direct inhibitory synaptic input mediated by specific interneuron subtypes. Elucidating these pathways will prove essential for understanding how feature-selective inhibition integrates into the broader cortical processing hierarchy. In this context, emerging functional connectomics datasets (e.g. Ding et al., 2025) promise invaluable in-sights by enabling researchers to bridge functional response profiles—including the dual-feature selectivity described here—with precise anatomical connectivity maps and cell-type-specific wiring motifs.

A further methodological consideration is that DreamSim was trained on human perceptual similarity judgments, while our neuronal data are from macaques. This cross-species application is supported by the deep homology between primate ventral visual streams: the anatomical and functional organization of V4 and the ventral pathway is broadly conserved between macaques and humans, and natural-image similarity judgments have been found to be highly consistent across the two species (e.g. **?**). Importantly, we deploy DreamSim not as a model of macaque perception but as an image-feature embedding to test whether stimuli that cluster in perceptual space evoke similar neuronal responses—a use that is robust to the precise calibration of the metric, as confirmed by control analyses (Fig. 9e,f). Nevertheless, a macaque-specific embedding would provide stronger grounding. We plan to address this point directly in our ongoing work: we are developing custom embeddings trained to align with macaque response geometry, using contrastive learning on MAI/LAI pairs from our own recordings, which will allow us to replace DreamSim with a representation grounded in the neural data itself.

Finally, we have not yet characterized the relationship between the most and least activating stimuli for individual neurons. Although our analyses identify images that elicit strong activation or suppression, we have not quantified the underlying tuning features—such as color, shape, texture, or their combinations—that give rise to these responses. Quantifying tuning along these feature dimensions will be essential for understanding how excitatory and suppressive stimuli are related. Qualitatively, our data suggest that there is no clear or consistent relationship between the two stimulus types; for example, we do not observe a systematic opponency in orientation or other visual dimensions. This observation aligns with Unk (2025), who similarly found that preferred and anti-preferred features appear to be independently distributed across neurons, with no apparent systematic relationship between the two feature types. More broadly, the concept of “opponency” itself requires clarification: does it reflect contrast along a single feature axis—such as high versus low spatial frequency—or coordinated shifts across multiple features, such as orientation, phase, and texture? Future work that integrates quantitative feature-tuning models with the functional selectivity profiles established here will be critical for determining whether excitatory and suppressive stimuli occupy distinct or systematically related regions within high-dimensional visual feature spaces.

### Normalization revisited: A role for feature-selective inhibition

Inhibitory cells in the brain exhibit a diversity of cell types comparable to that of excitatory neurons (Tasic et al., 2018; Gouwens et al., 2019; Yao et al., 2021), despite their much lower density (Keller et al., 2018). This raises the question: why is such diversity necessary? In the retina, a classic test bed for central brain research, the vast variety of inhibitory amacrine cells provides selective drive to distinct retinal ganglion cell types (e.g. Dia-mond, 2017; Matsumoto et al., 2025), shaping feature encoding and supporting parallel processing streams. Similarly, in the cortex, accumulating evidence indicates that inhibitory neurons target excitatory neurons with high specificity (Muñoz et al., 2017; Lu et al., 2017; Wu et al., 2023).

A recent millimetre-scale volumetric EM reconstruction of mouse visual cortex (Schneider-Mizell et al., 2025) mapped the connectivity of over 1,300 neurons, revealing that inhibitory neurons organize into motif groups with widespread target specificity. These motifs coordinate inhibition onto precise combinations of perisomatic and dendritic compartments across excitatory cell types, enabling compartment- and cell-type-specific modulation of cortical circuits far beyond broad class connectivity.

Likewise, dense connectomic reconstructions in the fly revealed that inhibitory interneurons, though few in number, constitute the majority of cell types and form highly specific, feature-selective connections to excitatory neurons (Matsliah et al., 2024; Sebastian Seung, 2024). For example, individual inhibitory cell types provide suppression to targeted single excitatory cell types at defined spatial scales, suggesting a division of labor across interneuron types. Sebastian Seung (2024) argues that such diversity is an inevitable consequence of implementing numerous highly specific normalization operations, paralleling the architectural principles seen in convolutional networks (see below).

Our results, together with the anatomical evidence discussed above, suggest that inhibition in sensory cortex is far more structured than traditionally assumed. Classical models treat cortical selectivity as driven by excitation to preferred features, with inhibition providing non-specific gain control via untuned normalization pools that sum across diverse feature preferences (Heeger, 1992; Carandini and Heeger, 2011).

In contrast, the dual-feature selectivity we describe may represent a biological implementation of specific rather than untuned normalization, where neurons apply suppressive filters to target specific anti-preferred features, rather than uniformly inhibiting all non-preferred inputs, thus enabling functions that extend beyond simple gain control. Indeed, Schwartz and Simoncelli (2001) demonstrated that non-uniform, feature-dependent normalization naturally emerges when the normalization weights are optimized to reduce statistical dependencies in natural signals, supporting the idea that such structured inhibition serves an efficient coding purpose. Related modeling work further shows that single-neuron activation functions—including those that permit negative responses—can themselves shape the circuit mechanisms and population geometries that emerge in recurrent networks (Tolmachev and Engel, 2025), suggesting that biological nonlinearities may likewise play a central role in organizing structured inhibition.

Structured suppression and normalization are also integral to modern artificial neural networks, albeit through different mechanisms. For example, batch normalization and attention mechanisms modulate activity based on the relationships between inputs (e.g. Ioffe and Szegedy, 2015; Ba et al., 2016; Vaswani et al., 2017). In our case, suppression is a fixed property of individual neurons: each neuron encodes the input by responding to preferred features and being suppressed by non-preferred ones. In contrast, artificial networks apply input-dependent modulation that is computed dynamically across inputs, such as through attention between tokens. Despite this difference, both artificial and biological networks appear to benefit from selectively attenuating certain features to improve representational efficiency.

Overall, our findings highlight feature-selective suppression as a general and underappreciated characteristic of cortical neurons. By defining specific feature dimensions through structured excitation and suppression, inhibition may enhance the representational capacity of individual neurons while preserving structured and interpretable response profiles that support flexible downstream readout—a principle that may be shared across sensory modalities and cognitive functions.

## ACKNOWLEDGEMENTS

The authors thank the International Max Planck Research School for Intelligent Systems (IMPRS-IS) for supporting PE. FHS is supported by the German Federal Ministry of Federal Ministry of Research, Technology and Space (BMFTR) via the Collaborative Research in Computational Neuroscience (CRCNS) (FKZ 01GQ2107), and FHS & KF are supported by the Collaborative Research Center (SFB 1233, Robust Vision, project number: 276693517). FHS and ASE acknowledges the support of the Lower Saxony Ministry of Science and Culture (MWK) with funds from the Volkswagen Foundation’s zukunft.niedersachsen program (project name: CAIMed Lower Saxony Center for Artificial Intelligence and Causal Methods in Medicine; grant number: ZN4257). EF acknowledges support from a European Research Council (ERC) grant (ERC-2022-STG, NEURACT, Grant agreement No: 101076710), the Hellenic Foundation for Research and Innovation (HFRI) under the 2nd Call for HFRI Research Projects to Support Faculty Members and Researchers with Grant Agreement No. 4049, and the HFRI under the “Funding of Basic Research (Horizontal support of all Sciences)” of the National Recovery and Resilience Plan “Greece 2.0” with funding from the European Union - NextGenerationEU with Grant Agreement No. 016552. MD acknowledges support from the European Union’s Horizon 2020 research and innovation program under the Marie Skłodowska-Curie Actions with Grant Agreement No. 101025482. ASE acknowledges support from the European Research Council (ERC) under the European Union’s Horizon Europe research and innovation programme (Grant agreement No. 101041669). KF acknowledges support from the European Research Council (ERC) under the European Union’s Horizon Europe research and innovation programme (Grant agreement No. 101117156). AST acknowledges support from the NIH (R01EY026927), the NSF (Collaborative Research in Computational Neuroscience, IIS-2113173), and the Defense Advanced Research Projects Agency (DARPA), Contract No. N66001-19-C-4020 and Contract No. DARPA NESD N66001-17-C-4002. The views, opinions and/or findings expressed are those of the author and should not be interpreted as representing the official views or policies of the Department of Defense or the U.S. Government. AST and SS acknowledge support from the Amaranth Foundation (Enigma Project).

## AUTHOR CONTRIBUTIONS NK

Conceptualization, Methodology, Software, Validation, Formal analysis, Investigation, Data Curation, Writing - Original Draft, Visualization, Project Administration **KF**: Conceptualization, Methodology, Software, Validation, Formal analysis, Investigation, Data Curation, Writing - Original Draft, Visualization, Project Administration, Funding Acquisition **KW**: Conceptualization, Methodology, Software, Data Curation **MD**: Investigation, Data Curation, Visualization, Writing - Review & Editing **KRa**: Conceptualization, Investigation, Methodology **PE**: Software **KRe**: Methodology, Investigation, Data Curation **PF**: Conceptualization, Data Curation **CN**: Methodology, Data Curation **TS**: Methodology, Data Curation **GG**: Methodology, Data Curation **SP**: Software, Methodology, Validation **AE**: Writing - Review & Editing **EYW**: Conceptualization, Data Curation **EF**: Writing - Review & Editing **SS**: Conceptualization, Funding Acquisition, Writing - Review & Editing **FHS**: Conceptualization, Supervision, Writing - Review & Editing, Funding Acquisition **AT**: Conceptualization, Supervision, Funding Acquisition, Writing - Review & Editing

## Materials and Methods

### Ethics and Animal Care

Data were collected from three healthy male rhesus macaques with approval from Baylor College of Medicine’s Institutional Animal Care and Use Committee (permit AN-4367). The monkeys were housed individually in a room with approximately ten other monkeys, allowing for rich social interactions on a 12-hour light/dark cycle. Regular veterinary care, balanced nutrition, and environmental enrichment were provided. All surgical procedures were performed under general anesthesia using aseptic techniques, with post-operative analgesics administered for seven days.

### Electrophysiological recordings

Non-chronic recordings were conducted using a 32-channel linear silicon probe (NeuroNexus V1*×* 32-Edge-10mm-60-177). Custom titanium recording chambers and head posts were surgically implanted. Prior to recordings, small trephinations (2 mm) were made over either (i) lateral V4, with eccentricities ranging from 1.7°to 18.3°of visual angle; or Medial V1, with eccentricities ranging from 1.4°to 3.0°of visual angle. A Narishige Microdrive (MO-97) and guide tube were used to carefully position the probes through the dura, taking care to minimize tissue compression.

Mice: Eight mice (*Mus musculus*: 4 male, 4 female) aged from 14 to 27 weeks were selected for experiments, with 2 females and 1 male expressing GCaMP6s in excitatory neurons via Slc17a7-Cre and Ai162 transgenic lines (stock nos. 023527 and 031562, respectively; The Jackson Laboratory) and the rest being C57BL/6J wildtype (stock no. 000664; The Jackson Laboratory). We performed acute recordings using Neuropixels probes 1.0 in awake, head-fixed mice according to (Jun et al., 2017). In brief, animals were implanted with a headpost and habituated to the experimental setup (head fixation on a treadmill) after recovery. On the recording day, the animals were briefly anaesthetized with isoflourane and a 1mm craniotomy was made above visual cortex (approximately 2.9 mm lateral to the midline sagittal suture and anterior to the lambda suture) (Froudarakis et al., 2014). The animals were then transferred to the experimental setup and allowed to recover from anaesthesia. Location of probe insertion was chosen according to stereotaxic coordinates for targeting V1, LM, and LI using Pinpoint (Birman et al., 2023), with all penetrations ranging from 600–1100 *µm* on the anteroposterior axis, 2900–3500 *µm* on the mediolateral axis, and at an angle of 55°or 60°with respect to the ventrodorsal axis. One probe was smoothly lowered through the craniotomy to the final depth according to the trajectory planning with Pinpoint (Birman et al., 2023) to cover the whole cortex (covering 1800–2000 *µm* of the probe) and allowed to settle for approximately 20 minutes before any recording.

### Data collection and processing

Electrophysiological data was recorded as a broadband signal (0.5Hz-16kHz) and digitized at 24 bits. For spike sorting, the 32-channel array was divided into 14 groups of six adjacent channels. Spikes were detected when signals exceeded five times the standard deviation of noise. Principal component analysis was used for feature extraction, and a Kalman filter mixture model tracked waveform drift. Single-unit isolation was manually verified by assessing stability, refractory periods, and principal component plots.

For mice, neuronal activity recordings were made with custom-written software in LabView and then automatically spike sorted with the Kilosort3 spike sorting software (Pachitariu et al., 2023). Neurons automatically classified as “single units” and that showed reliable firing during visual stimuli presentation were used for the model, in total: 598 V1, 350 LM, and 126 LI neurons from 20 recording sessions.

### Visual stimulation

Stimuli were displayed on a 23.8” LCD monitor (100 Hz refresh rate, 1920 *×* 1080 resolution) positioned 100 cm from the subjects (*≈* 63 pixels/degree). A camera-based eye tracking system verified that monkeys maintained fixation within *≈* 0.95° of a small red fixation target. After maintaining fixation for 300 ms, visual stimuli were presented. Successful fixation throughout the trial resulted in a juice reward. For mice, natural images were presented 15 cm away from the left eye with a 23.8” LCD monitor (100 Hz refresh rate, 1920 *×* 1080 resolution). We positioned the monitor so that it was centered on and perpendicular to the surface of the eye at the closest point, corresponding to a visual angle of 2.2°/cm on the monitor.

### Receptive field mapping & stimulus placement

Receptive fields were mapped at the beginning of each session using a sparse random dot stimulus. A single dot (0.12-1° in size) was displayed on a gray background, changing position and color (black or white) every 30 ms during two-second fixation trials. Multi-unit receptive field profiles were obtained through reverse correlation, and population receptive fields were estimated by fitting a 2D Gaussian to the spike-triggered average.

For V1 recordings, the fixation spot was kept at the center of the screen, and grayscale natural image stimuli (6.7°in size) were centered at the mean receptive field location. The remainder of the screen was kept gray. For V4 recordings, color natural image stimuli covered the entire screen, and the fixation spot was positioned to place the mean receptive field as close as possible to the screen’s center. Due to recording site locations, this typically placed the fixation spot near the upper left border of the screen.

For mice: Visual area segmentation was performed by mapping the reversals of the retinotopy based on the RF progression along the probe as described previously (Tafazoli et al., 2017).

### Stimulus selection

We selected 24, 075 images from 964 ImageNet categories, cropped to 420 *×* 420 pixels with 8-bit intensity resolution. From this set, 75 images were designated as the test set, 20% of the remaining images as the validation set, and the rest (19, 200 images) as the training set. The same image sets were used for both V1 and V4 recordings, with some differences in preprocessing (see below).

For a subset of V4 experiments, the dataset was augmented with rendered scenes, resulting in an equal mix of ImageNet and rendered images in the training, validation, and test sets. This synthetic dataset of rendered 3D scenes was created using Kubric (Greff et al., 2022) and Blender ((Blender Foundation, 2024)). We first manually created 10 primitive 3D objects in Blender, including basic geometric shapes (spheres, cubes, cylinders, cones, pyramids, etc.) and simple composite forms. Each object was exported as a Wavefront OBJ file with accompanying material (MTL) and texture coordinate information to ensure consistent UV mapping across all renders. For surface textures, we utilized the Describable Textures Dataset (Cimpoi et al., 2014), which contains 47 texture categories with diverse visual properties ranging from regular patterns (e.g., striped, checkered) to stochastic textures (e.g., marbled, bubbly). We developed an automated rendering pipeline that generated 200, 000 unique scenes at 420 *×* 236 pixel resolution, matching the aspect ratio used in our V4 experimental stimuli. Each rendered scene consisted of a single 3D object with UV-mapped texture randomly sampled from the DTD dataset, placed against a background with a different randomly selected DTD texture. To ensure comprehensive sampling of the visual parameter space, we systematically varied multiple scene attributes: (1) object identity (uniformly sampled from the 10 primitive shapes), (2) object position (*x, y, z* coordinates sampled within the camera frustum), (3) orientation (random rotation quaternions), (4) scale, and (5) lighting conditions (directional light with varying intensity and angle). Shadow rendering was enabled to introduce natural occlusion patterns and enhance depth cues. This systematic variation ensured that our synthetic dataset captured a broad range of feature combinations while maintaining precise control over individual visual attributes.

### Stimulus presentation

Each trial involved 2.1 seconds of continuous fixation, including 300 ms of gray screen at the beginning and 15 consecutive images displayed for 120 ms each without gaps. Test images were repeated 20-50 times throughout the session, while training and validation images were shown only once. The animal completed up to 1400 trials per day, resulting in up to 20000 stimulus-response pairs per neuron.

For V1 recordings, grayscale images were displayed at their original resolution covering a 6.7° visual angle, with the fixation spot at the center of the screen and the images centered at the mean receptive field location. The rest of the screen remained gray. Neural responses were analyzed within a 40-160 ms window following stimulus onset. In contrast, for V4 recordings, full-color images were upscaled to match the screen width while maintaining their aspect ratio, with upper and lower bands cropped to fill the entire screen. The fixation spot was positioned to place the mean receptive field as close as possible to the screen’s center, typically near the upper left border due to recording site locations. Spike counts for V4 were collected within a 60-160 ms window after stimulus onset.

For mice, 5, 100 natural images from ImageNet (ILSVRC2012) were cropped to fit a 16 : 9 monitor aspect ratio and converted to gray scale. To collect data for training a predictive model of the brain, we showed 5, 000 unique images as well as 100 additional images repeated 10 times each. This set of 100 images were shown in every recordings for evaluating cell response reliability within and between recordings. Each image was presented on the monitor for 500 ms followed by a blank screen lasting between 100 and 200 ms, sampled uniformly.

### Image pre-processing

We employed two distinct image processing pipelines to prepare stimuli for model training and evaluation. In the V4 pipeline, starting with original images of 420 *×* 420 pixels at a resolution of 14 px*/*^*°*^, we cropped the upper and lower bands to fit the full screen for presentation, resulting in images of 420 *×* 236 pixels. We then extracted only the bottom center 200 *×* 200 pixel region because of the location of the receptive fields (RFs) and subsequently downsampled these images to 100 *×* 100 pixels (corresponding to either 5.8 px*/*^*°*^ or 7 px*/*^*°*^) for model training. For the V1 approach, we cropped the central 2.65^*°*^ (167 pixels) of the original 420 *×* 420 image at its resolution of 63 px*/*^*°*^ and applied bicubic interpolation to downsampled to 93 *×* 93.

### Model architectures & training

For macaque V1 data, following Fu et al. (2024), we used a ConvNext-v2-tiny (Woo et al., 2023) architecture as the core, with original weights from the huggingface transformers library. After hyperparameter search, we selected the *stages-1-layers-0* as the optimal output layer. We applied a Gaussian read-out approach (Lurz et al., 2022) to transform the core feature maps into neuronal responses. This readout learns the coordinates of each neuron’s receptive field center on the feature maps and implements a 2D isotropic Gaussian distribution to extract features from this location. During training, the readout samples positions according to this distribution, gradually focusing on the optimal receptive field location, while at inference time it uses the learned fixed positions. The extracted features were then processed through a neuron-specific affine projection with ELU non-linearity to predict the scalar neuronal activity.

For V1 model training, following (Fu et al., 2024), we minimized the Poisson loss between recorded and predicted neuronal activity. We first trained the readout for 20 epochs with frozen core weights, then reduced the learning rate from 0.001 to 0.0001 and optimized both the ConvNext core and readout weights using the AdamW optimizer (Loshchilov and Hutter, 2017) for 200 epochs. We trained an ensemble of five models with different random seeds and used their averaged predictions for all analyses.

Our neural predictive model of primate V4 consisted of a pretrained *core* computing nonlinear features from input images, and a *Gaussian readout* (Lurz et al., 2022) mapping these features to single neuron responses. Following (Willeke et al., 2023), we used an adversarially trained ResNet50 (Salman et al., 2019) as the core, with the first residual block of layer 3 (layer3.0) providing the feature maps, as this configuration yielded the highest predictive performance. The parameters of this pretrained network remained fixed during training. After batch normalization and ReLU activation, we obtained a nonlinear feature space shared across all neurons. For each V4 neuron, the Gaussian readout learned the receptive field center position on the output tensor to extract a feature vector. The readout implemented a 2D isotropic Gaussian distribution, sampling locations during training and using fixed positions during inference. The extracted features were then processed through a linear-nonlinear model with *L*_1_-regularized weights and an ELU+1 nonlinearity (Clevert et al., 2015) to ensure positive responses.

For the V4 model training, as in (Willeke et al., 2023), we minimized the summed Poisson loss across neurons between observed and predicted spike counts, with added *L*_1_ regularization on the readout parameters. During training, we zeroed gradients for neurons not shown a particular image. We used a batch size of 64, the Adam optimizer (Kingma and Ba, 2014) with an initial learning rate of 3 · 10^*−*4^ and momentum of 0.1. We implemented early stopping based on validation loss, decaying the learning rate by 0.3 after five epochs without improvement, and stopping after four such decay steps. We trained an ensemble of five models with different random seeds and used their averaged predictions for all analyses.

For the mouse model, we trained a digital twin model using the Sensorium competition model architecture (Willeke et al., 2022) with neural data recorded across V1 (*n* = 598 neurons), LM (*n* = 350 neurons), and LI (*n* = 126 neurons). Specifically, a convolutional core shared across all neurons from all areas combined with neuron-specific readouts, trained end-to-end to predict neuronal responses to natural images.

### Model evaluation

To evaluate both V1 and V4 models, we measured performance using correlation to average (Franke et al., 2022; Cadena et al., 2023; Willeke et al., 2022) on held-out test images. This metric computes the correlation between model predictions and the average neuronal responses across repeated presentations of the same stimuli. By comparing predictions to trial-averaged responses rather than single-trial responses, this approach focuses on how well the models capture the stimulus-driven component of neural activity while accounting for biological trial-to-trial variability. Following (Fu et al., 2024) for V1 and (Willeke et al., 2023) for V4, we applied this evaluation consistently across both areas to enable fair comparison of model performance. Models trained exclusively on ImageNet generalized well to rendered images, showing no significant differences in prediction performance (data not shown). Consequently, we refer to both datasets collectively as ‘naturalistic images’ in the Results section. Detailed information regarding which dataset was used for each analysis is provided in the figure legends and the Methods section below.

### Identification of most and least activating images

To identify naturalistic images that elicited extreme responses from modeled neurons, we conducted a large-scale screening across two complementary sources: 200, 000 synthetically rendered scenes with controlled shape and texture variations, and 1, 281, 167 natural images from the ImageNet-1K training set (Deng et al., 2009). Prior to neuronal response prediction, all images underwent standardized preprocessing to match the experimental conditions. Images were center-masked using the mean receptive field profile computed from the population of synthesized most and least exciting images for V1 and V4 neurons, respectively. This masking procedure ensured that visual features were evaluated within the approximated retinotopic locations while controlling the influence of peripheral image regions. Additionally, we normalized all images to fixed *ℓ*_2_ norms (12.0 for V1, 40.0 for V4) to control for overall contrast differences and ensure fair comparison across diverse image content. These normalization values were empirically determined to match the typical contrast range of natural images while preventing saturation artifacts. For response prediction, we employed area-specific models: the ConvNeXt-based architecture for V1 neurons and the adversarially-trained ResNet50 for V4 neurons, as described in the model architecture section. Each model computed predicted firing rates for all neurons across both image datasets, resulting in response matrices of 200, 000Ö*N* and 1, 281, 167Ö*N* for rendered and natural images respectively, where *N* represents the number of recorded neurons. From these combined predictions, ranked in ascending order per neuron, we identified the top and bottom images for each neuron, corresponding to the most and least activating images (MAIs and LAIs). Similarly for mice, screening was conducted across the 200, 000 synthetically rendered scenes and images were normalized to a fixed *ℓ*_2_ norm (10.0 for V1, LM, and LI). For response prediction we employed the one model for all three areas, as described in the model architecture section. The model computed predicted firing rates for all neurons in the dataset, resulting in a response matrix of 200, 000Ö*N*, where *N* represents the total number of recorded neurons recorded across V1, LM, and LI (*N* = 1074).

### Optimization of most and least exciting images

We employed gradient-based optimization in the pixel space (for V1 neurons) or frequency (for V4 neurons) domain to generate images that optimally drove or suppressed individual neuronal responses (Fel et al., 2023). Starting from random noise images, we iteratively modified pixel values to either maximize (for most exciting inputs, MEIs) or minimize (for least exciting inputs, LEIs) the predicted neuronal response. For V4, this optimization operated exclusively on the phase spectrum of the Fourier transform while constraining the amplitude spectrum to match a fixed mean amplitude computed over 10, 000 randomly sampled ImageNet images, ensuring that synthesized images maintained realistic spatial frequency content. For V1, the image synthesis was achieved by directly modifying the image pixel values themselves. During optimization, we applied *ℓ*_2_ norm constraints matching those used in the screening procedure (12.0 for V1, 40.0 for V4) to maintain consistent contrast levels across all analyses. To improve optimization robustness and avoid local minima, we implemented a multi-crop augmentation strategy. This operation generated 4 random crops per image at each optimization step, with crop centers sampled from a Gaussian distribution (*µ* = 0.5, *σ* = 0.15) relative to image dimensions. Importantly, we maintained a fixed box size of 1.0 throughout optimization, meaning crops spanned the full image extent with only positional jittering. This approach effectively provided the optimizer with multiple gradient estimates per iteration by evaluating slightly shifted versions of the image, similar to translation data augmentation but applied during the optimization process itself. The crop sizes included additional Gaussian noise (*σ* = 0.05) clamped between 0.05 and 1.0, though with our box size fixed at 1.0, this primarily introduced minor scale variations around the full image size. Each crop was resized back to the original dimensions (93 *×* 93 pixels for V1, 100 *×* 100 pixels for V4) using bilinear interpolation before neural response prediction. By averaging gradients across these multiple crops, the optimization procedure became more robust to small spatial shifts, encouraging the emergence of features that drove consistent neuronal responses. We performed 256 optimization steps using the Adam optimizer ((Kingma and Ba, 2017)) with a learning rate of 0.05, generating both MEIs and LEIs for each recorded neuron.

### Verification of LAIs & MAIs and LEIs & MEIs

To verify that model-identified least and most activating stimuli accurately represented neuronal preferences, we implemented two complementary validation approaches. To validate predictions against in vivo recordings, we selected the single images from held-out test sets predicted to elicit the strongest and weakest responses for each neuron. We then determined where these images ranked within each neuron’s recorded response distribution, allowing us to assess whether model predictions aligned with actual neuronal behavior.

To test generalization across models, we trained independent evaluator models. For V1 neurons, we used the same ConvNeXt architecture as our generator model but with independent initialization, fine-tuning the core network while training neuron-specific readouts from scratch. For V4 neurons, we employed an entirely different architecture—an attention-based convolutional network trained end-to-end. A distinct architecture was necessary here because the V4 generator model uses a fixed, pretrained ResNet50 backbone whose weights are deterministic: any re-trained model sharing this backbone would not constitute a genuinely independent evaluation. By contrast, for V1, the ConvNeXt core is fine-tuned from different random initializations, producing architecturally equivalent but computationally independent models. To evaluate stimulus effectiveness, we presented each evaluator model with the four synthetic images (MEI, LEI, MAI, LAI) derived from the generator model for each neuron. To contextualize response strength, we compared these against a reference set of 200, 000 contrast- and size-matched naturalistic images. For each synthetic image, we calculated its response percentile, defined as the proportion of reference images evoking a weaker model-predicted response. This cross-model validation allowed us to determine whether identified stimuli represented genuine neuronal tuning properties rather than model-specific artifacts. The choice of distinct architectures for V1 and V4 evaluator models reflects a difference in how each generator is trained: the V1 generator fine-tunes its ConvNeXt core, yielding genuine independence across random seeds, whereas the V4 generator relies on a fixed pretrained ResNet50 backbone shared identically across all instances. A truly independent V4 evaluator therefore required a fundamentally different architecture.

### Baseline firing rate

To extract baseline firing rates, we counted spikes during the 300 ms fixation window prior to stimulus onset. We converted these counts into firing rates (Hz) and computed the mean baseline activity across all valid trials. During this fixation period, the screen was set to mean gray level (127 in 8-bit). For mice the baseline firing rates (Hz) were calculated during the blank screen (200ms duration) before the stimulus onset.

### DreamSim image similarity

To test whether least and most activating rendered images (LAIs and MAIs) contain robust feature combinations and exhibit similarity within categories, we embedded all rendered images into the image similarity model DreamSim (Fu et al., 2023b). DreamSim leverages deep neural network representations to quantify perceptual image similarity in a manner that correlates with human judgments. We used the penultimate layer as an embedding for each image. For V1, we preprocessed images by masking them with the mean receptive field mask of V1 neurons, normalizing them to have the same contrast, and converting them to grayscale. For V4, we masked images with the V1 mask, normalized them, and preserved their color information. To quantify similarity, we computed pairwise cosine similarity between DreamSim vectors for the top and bottom 10 images per neuron (within-MAI and within-LAI comparisons). Additionally, we calculated pairwise distances between MAIs and LAIs, as well as distances from both MAIs and LAIs to 10 randomly selected images. This analysis resulted in cosine similarity distributions per neuron and condition. We used d-prime as a measure of discriminability, with values below 0.5 indicating low discriminability between image categories.

### Population-level analysis

To assess the extent to which the most activating images (MAIs) and least activating images (LAIs) of a given neuron also activate other neurons within the same area, we performed the following analysis (for both species). For each neuron, we first identified its top 15 MAIs and bottom 15 LAIs. We then evaluated how these images ranked in terms of activation for every other neuron in the population. Specifically, for each other neuron, we computed the percentile rank of each MAI and LAI within its response distribution obtained from the entire set of size- and contrast-matched naturalistic images (*>* 1 million images). This yielded a distribution of percentile ranks for the MAIs (and LAIs) of each neuron across all other neurons. Percentile ranks were then averaged across the 15 MAIs and 15 LAIs for each neuron. As a control, we randomly sampled 15 images from the full set of naturalistic images (i.e. *>* 1 million images) and computed their percentile ranks in the same way. Additionally, to examine whether this effect generalizes across individuals, we conducted a cross-monkey analysis. For this, we identified the MAIs and LAIs of neurons recorded in one monkey and calculated the response percentiles of these images for neurons recorded in the other monkey.

### Data and code availability

Code for screening images in the digital twin, optimizing images, and using the digital twin is available at: https://github.com/enigma-brain/dualneuron. Data, including predicted responses to ImageNet images, rendered images, as well as code to load ImageNet images, will be available at: DOI:10.5061/dryad.q573n5tx3.

## Supplementary Information

**Supplemental Fig. 1.**
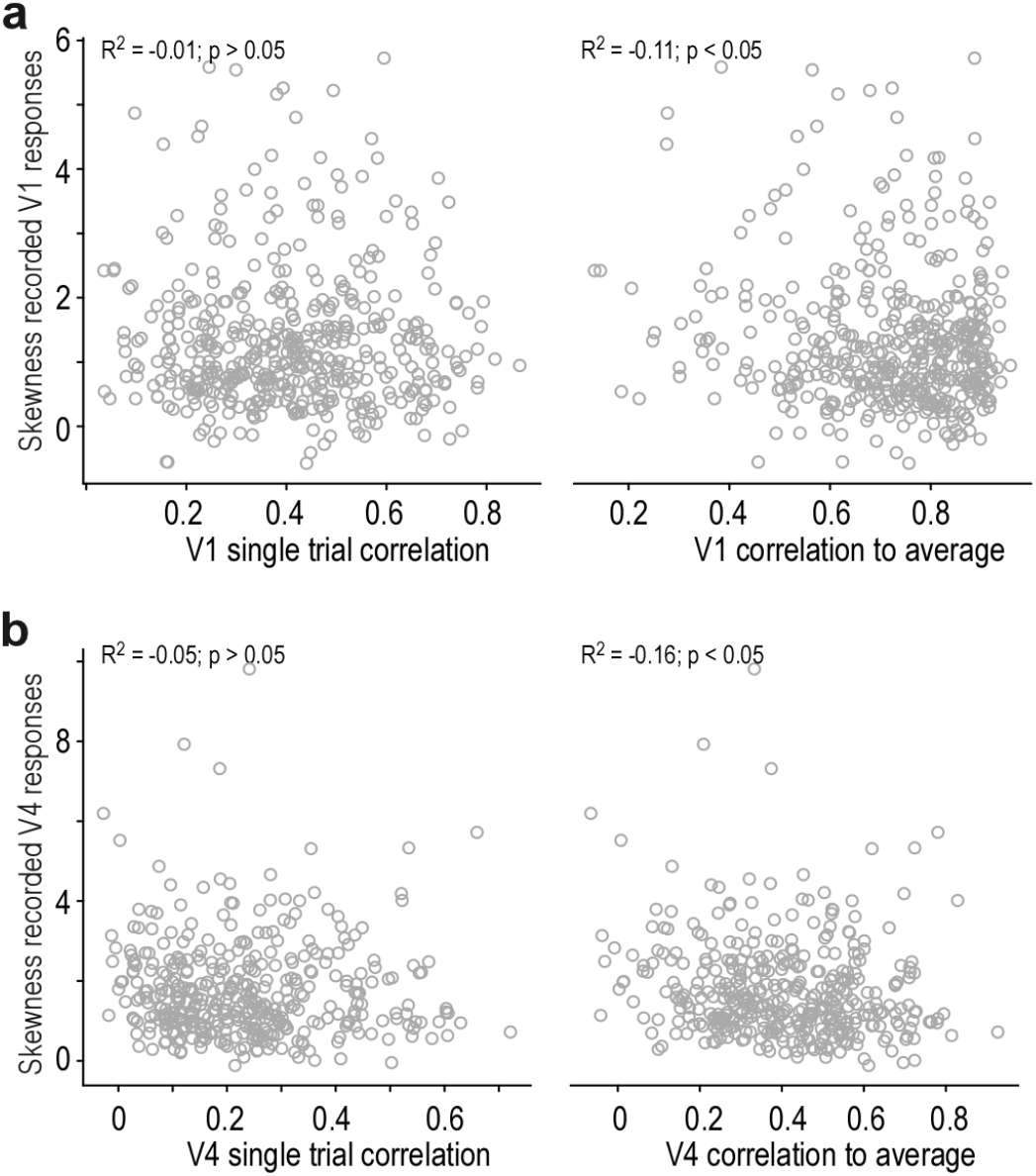
Relationship between model performance and response skewness. **a**, Single trial correlation (left) and correlation to average (right) of V1 model plotted versus skewness of recorded V1 responses. **b**, Like (a), but for V4 model and responses.

**Supplemental Fig. 2.**
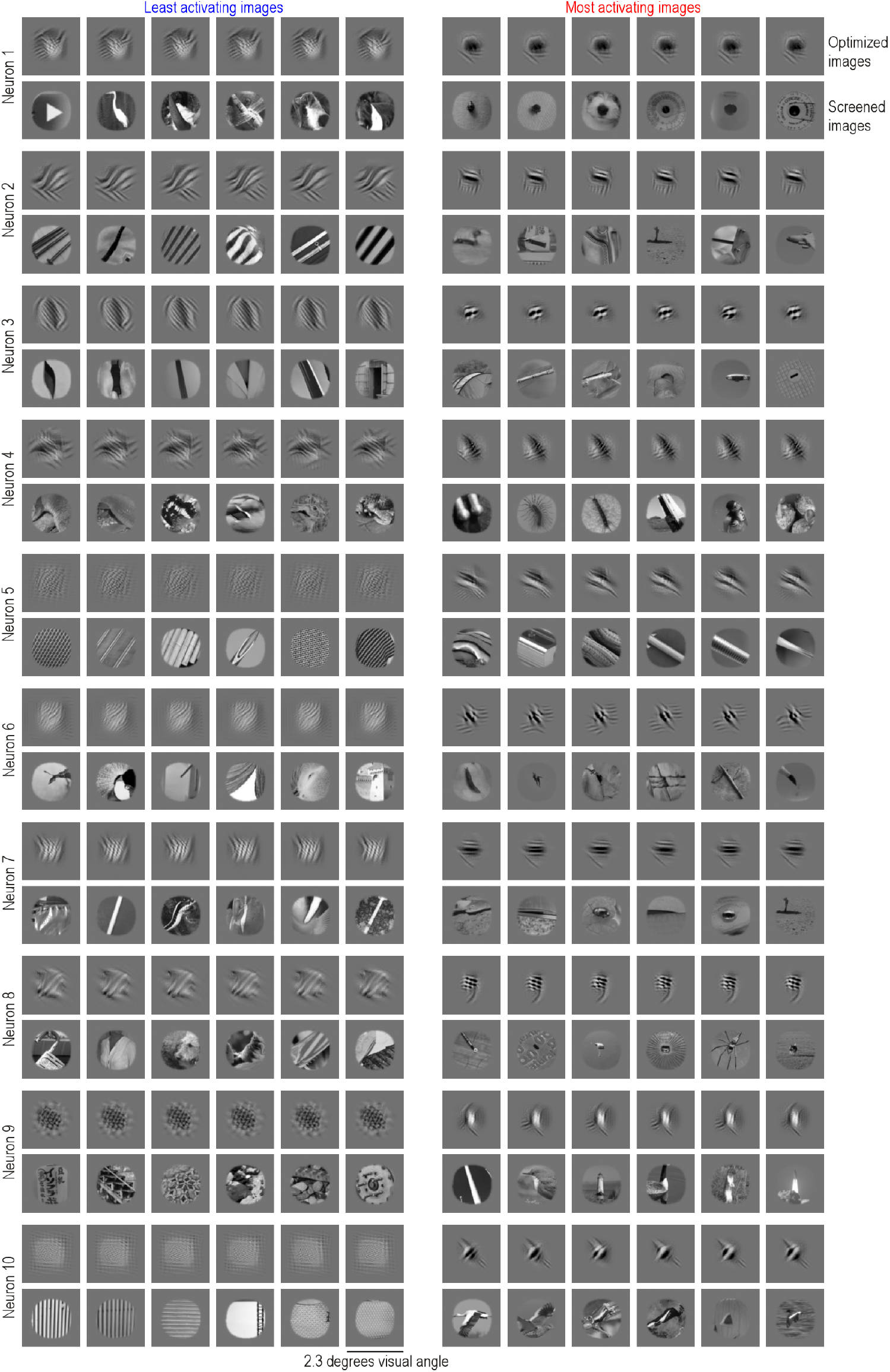
Most and least activating images of example V1 neurons. Least activating (left) and most activating (right) images of ten example V1 neurons. Per neuron, the top row shows optimized images (LEIs and MEIs) and the bottom row shows screened ImageNet images (LAIs and MAIs). Each image is 2.3 *×* 2.3 degrees visual angle, with the receptive field of the neuron in the center.

**Supplemental Fig. 3.**
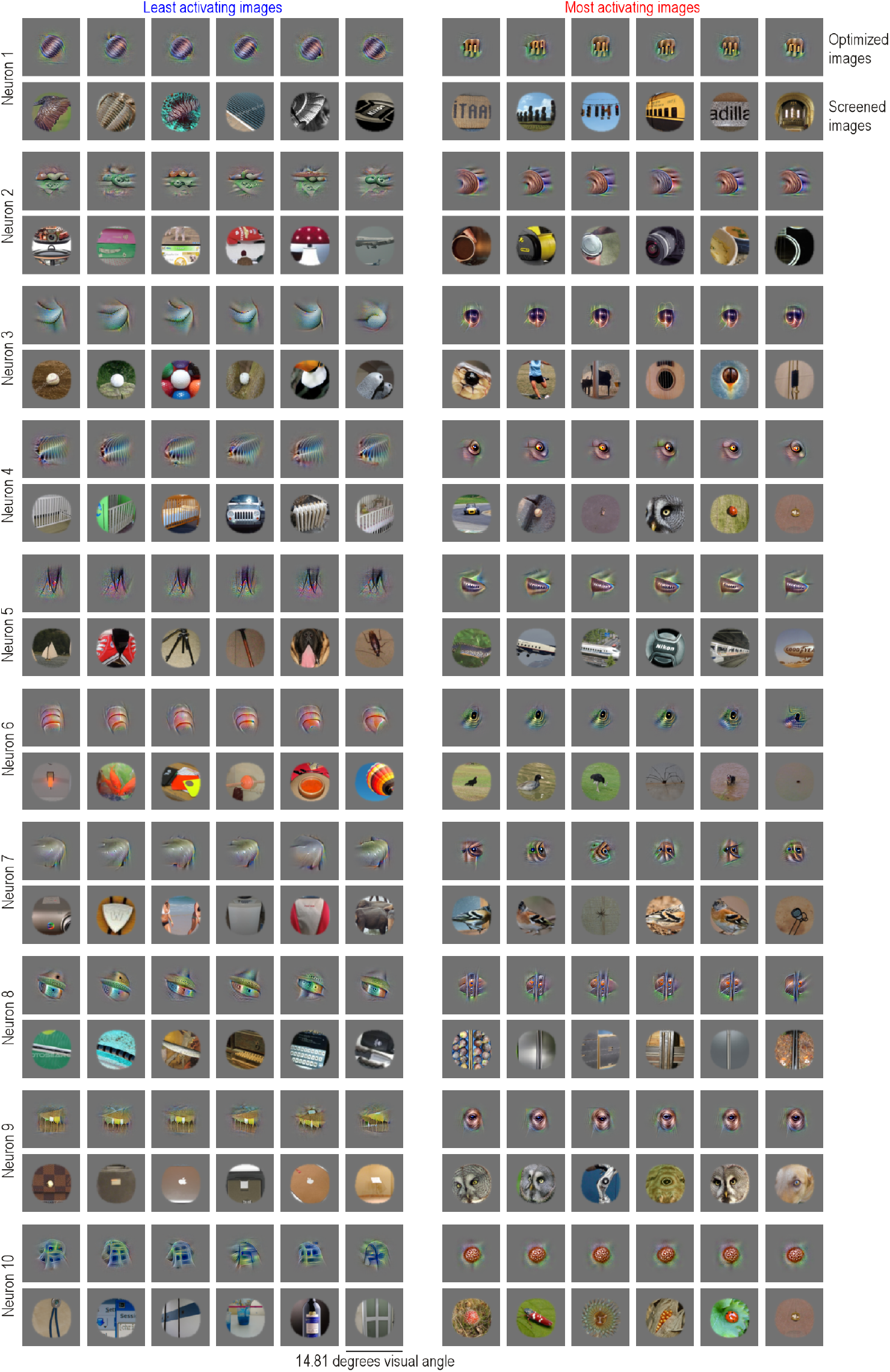
Most and Least Activating Images of Example V4 Neurons. Least activating (left) and most activating (right) images of ten example V4 neurons. Per neuron, the top row shows optimized images (LEIs and MEIs) and the bottom row shows screened ImageNet images (LAIs and MAIs). Each image is 14.83 *×* 14.83 degrees visual angle, with the receptive field of the neuron in the center.

**Supplemental Fig. 4.**
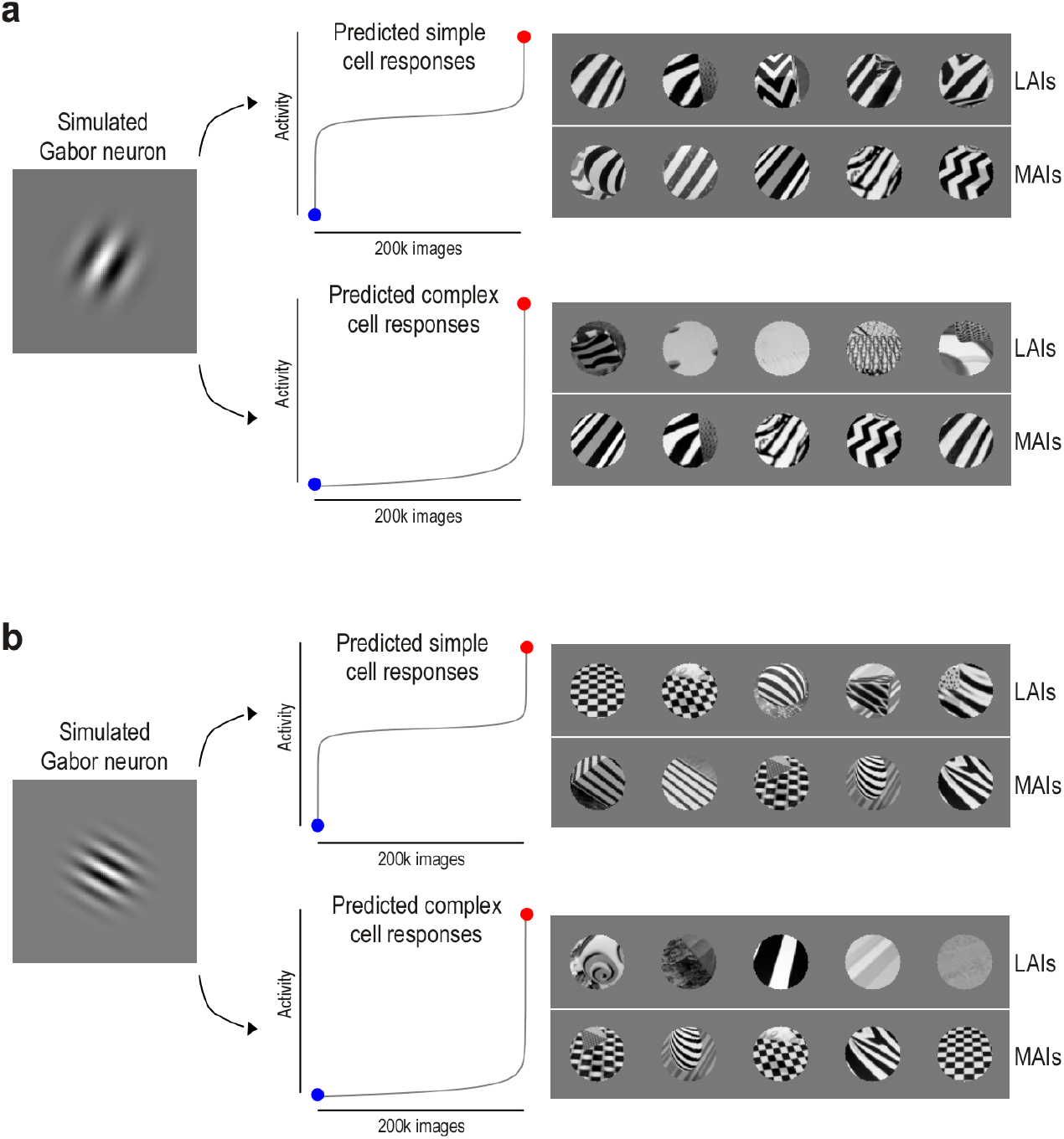
Most and least activating images of simulated simple and complex cells. **a**, Example simulation of a V1 simple and complex cell based on the same Gabor receptive field (left). Activations to 200, 000 naturalistic images were computed as the dot product between the Gabor filter and each image. For simple cells, this revealed least (blue) and most (red) activating images, showing a bimodal response distribution with LAIs sharing the same orientation but differing in phase. Complex cells were modeled by squaring and pooling responses across different phases, resulting in sparse activation profiles without coherent LAIs.

**Supplemental Fig. 5.**
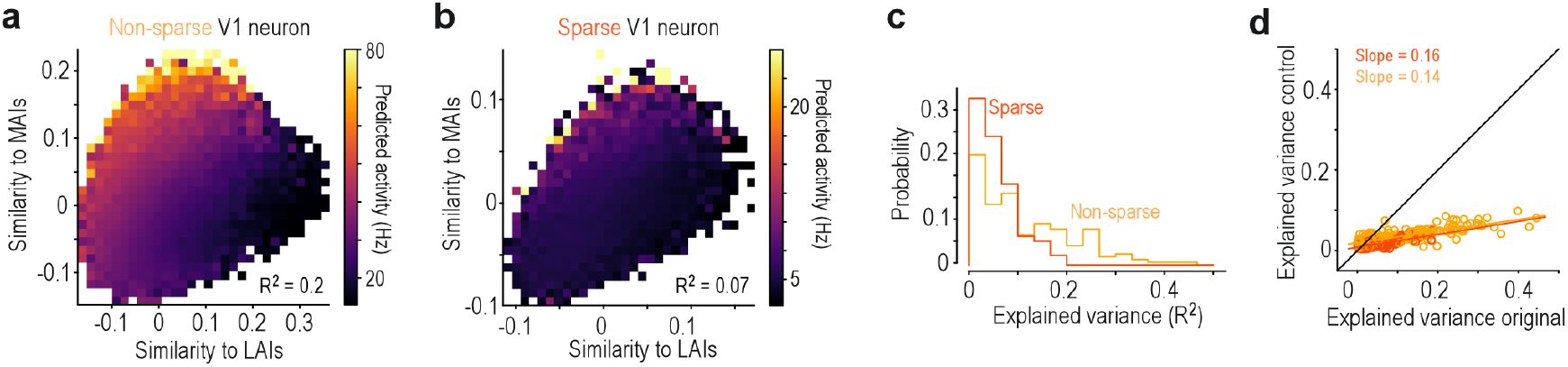
Responses of V1 neurons vary continuously depending on preferred and non-preferred stimuli. **a**, Example 2D DreamSim similarity space (cf. Fig. 9) for a non-sparse V1 neuron. Bins are color-coded by the mean predicted neuronal activity of the images within each bin. The *R*^2^ indicates the variance explained by a linear fit. The color bar spans the 0.1 to 99.9th percentile of neuronal responses across all images. **b**, Same as (b), but for a sparse V1 neuron. Here, the activity gradient aligns primarily along the *y*-axis, indicating stronger modulation by similarity to the MAI than to the LAI. **c**, Variance explained (*R*^2^) by linear regression predicting neuronal activity from the 2D similarity space (as in a,b), shown separately for sparse (dark orange) and non-sparse (light orange) V4 neurons. The explained variances values are generally lower for V1 compared to V4 neurons (cf. Fig. 9). This is likely due to the fact that the DreamSim representational space is more closely aligned with the representational space of V4 than V1. **f**, Variance explained (*R*^2^) by linear regression for the original analysis (similarity to MAIs and LAIs on *x*- and *y*-axes) and for Control 1 (left), where both MAIs and LAIs were replaced with random images.

